# A cathepsin C-like protease post-translationally modifies *Toxoplasma gondii* secretory proteins for optimal invasion and egress

**DOI:** 10.1101/2023.01.21.525043

**Authors:** L. Brock Thornton, Melanie Key, Chiara Micchelli, Andrew J. Stasic, Samuel Kwain, Katherine Floyd, Silvia N. J. Moreno, Brian N. Dominy, Daniel C. Whitehead, Zhicheng Dou

## Abstract

Microbial pathogens use proteases for their infections, such as digestion of proteins for nutrients and activation of their virulence factors. As an obligate intracellular parasite, *Toxoplasma gondii* must invade host cells to establish its intracellular propagation. To facilitate invasion, the parasites secrete invasion effectors from microneme and rhoptry, two unique organelles in apicomplexans. Previous work has shown that some micronemal invasion effectors experience a series of proteolytic cleavages within the parasite’s secretion pathway for maturation, such as the aspartyl protease (TgASP3) and the cathepsin L-like protease (TgCPL), localized within the post-Golgi compartment (1) and the endolysosomal system (2), respectively. Furthermore, it has been shown that the precise maturation of micronemal effectors is critical for *Toxoplasma* invasion and egress (1). Here, we show that an endosome-like compartment (ELC)-residing cathepsin C-like protease (TgCPC1) mediates the final trimming of some micronemal effectors, and its loss further results in defects in the steps of invasion, egress, and migration throughout the parasite’s lytic cycle. Notably, the deletion of TgCPC1 completely blocks the activation of subtilisin-like protease 1 (TgSUB1) in the parasites, which globally impairs the surface-trimming of many key micronemal invasion and egress effectors. Additionally, we found that TgCPC1 was not efficiently inhibited by the chemical inhibitor targeting its malarial ortholog, suggesting that these cathepsin C-like orthologs are structurally different within the apicomplexan phylum. Taken together, our findings identify a novel function of TgCPC1 in the processing of micronemal proteins within the secretory pathway of *Toxoplasma* parasites and expand the understanding of the roles of cathepsin C protease.

**IMPORTANCE:** *Toxoplasma gondii* is a microbial pathogen that is well adapted for disseminating infections. It can infect virtually all warm-blooded animals. Approximately one-third of the human population carries toxoplasmosis. During infection, the parasites sequentially secrete protein effectors from the microneme, rhoptry, and dense granule, three organelles exclusively found in apicomplexan parasites, to help establish their lytic cycle. Proteolytic cleavage of these secretory proteins is required for the parasite’s optimal function. Previous work has revealed that two proteases residing within the parasite’s secretory pathway cleave micronemal and rhoptry proteins, which mediate parasite invasion and egress. Here, we demonstrate that a cathepsin C-like protease (TgCPC1) is involved in processing several invasion and egress effectors. The genetic deletion of *TgCPC1* prevented the complete maturation of some effectors in the parasites. Strikingly, the deletion led to a full inactivation of one surface-anchored protease, which globally impaired the trimming of some key micronemal proteins before secretion. Therefore, this finding represents a novel post-translational mechanism for the processing of virulence factors within microbial pathogens.

## INTRODUCTION

*Toxoplasma gondii*, a eukaryotic pathogen belonging to the Apicomplexa phylum, widely spreads its infection in virtually all warm-blooded animals, including humans (3, 4). During infection, the parasites penetrate and hijack the host’s plasma membrane to form their own niche within the host cells for intracellular replication. Upon exhausting the nutrients from host cells, the parasites egress to pursue new hosts. Proteases play crucial roles throughout the individual steps within the lytic cycle of *Toxoplasma* parasites, such as TgASP3, which is localized in the post-Golgi apparatus and mediates the maturation of microneme and rhoptry proteins for parasites invasion (1). Additionally, TgSUB1, a GPI-anchored serine protease, processes microneme proteins at the parasite’s surface for invasion and egress (5).

Genome annotation has revealed that *Toxoplasma* parasites encode hundreds of proteases (6). Previous findings reported that the parasites possess a lysosome-like acidic organelle, named the plant-like vacuolar compartment (PLVAC) (7). The acidic hydrolases stored in this organelle are used to maturate some micronemal proteins and digest ingested host proteins, which facilitate parasite invasion and replication (2, 8–10). A few orthologs of classic lysosomal proteases have been identified in the PLVAC, such as cathepsin L (TgCPL), cathepsin B (TgCPB), and one cathepsin D-like (TgASP1) proteases in the PLVAC (9–11). Among these proteases, TgCPL is a master protease that mediates the maturation of TgCPB and TgASP1 (10, 11). The loss of TgCPL results in defective invasion and growth in tachyzoites and reduced acute and chronic virulence (9, 12). Within both acute and chronic infections, mutants lacking *TgCPB* or *TgASP1* did not display any growth defects nor virulence loss (10, 11). To maintain optimal proteolytic activities within the PLVAC, the parasites express two transmembrane proton pumps for luminal acidification of the organelle, termed the plant-like pyrophosphatase (TgVP1) and the vacuolar ATPase complex (v-ATPase) (13, 14). The *TgVP1*-deletion mutant is viable and displays defective microneme secretion, invasion, and reduced extracellular survival (13). The mutant containing a non-functional v-ATPase does not maturate microneme and rhoptry proteins properly, further compromising the parasite’s lytic cycle and virulence (14). Therefore, proteases within the parasite’s endolysosomal pathway play a key role in parasite infections.

Cathepsin C protease, an aminopeptidase, is located in the lysosome in many eukaryotic organisms (15, 16). The mammalian cathepsin C, also known as dipeptidyl peptidase I (DPP-I), is involved in the activation of other proteases, such as neutrophil elastase, cathepsin G, neutrophil serine protease 4 (NSP4), and granzymes A and B (16). *Toxoplasma* encodes three cathepsin C-like proteases (17) as shown by an ortholog-based genome annotation (www.toxodb.org), named cathepsin C-like protease 1, 2, and 3 (TgCPC1, TGGT1_289620; TgCPC2, TGGT1_276130; TgCPC3, TGGT1_267490). *Plasmodium spp*., closely related to *Toxoplasma*, also express 3 cathepsin C orthologs, named dipeptidyl aminopeptidases (PfDPAP1-3) (18, 19). Malarial PfDPAP1 and 3 (PF3D7_1116700 and PF3D7_0404700, respectively) were localized to the digestive vacuole, an organelle equivalent to the PLVAC, for digestion of incorporated hemoglobins. PfDPAP1 is also observed in the parasitophorous vacuole (PV) (20). PfDPAP2 is a gametocyte-specific gene, and its function still remains unclear. Both PfDPAP1 and PfDPAP3 are essential for the pathogenesis of malaria parasites (20–22). In contrast to malaria parasites, TgCPC1 and TgCPC2 were reported as dense granule proteins localized to the PV (17), while TgCPC3 is exclusively expressed in the sporozoite stage (17). A previous study successfully deleted *TgCPC1* in the parasites, suggesting that TgCPC1 is dispensable during *Toxoplasma* infections. The primary structure analysis revealed that both TgCPC1 and TgCPC2 contain signal peptides at their N-termini, implying that they traffic through the parasite’s endolysosomal system. The latest publication characterizing the subcellular proteomic atlas of *Toxoplasma* found TgCPC1 in the micronemes (23), also indicating its access to the endolysosomal system. These discrepancies prompted our re-evaluation of the roles of TgCPC1 in *Toxoplasma* infections.

Here, we reveal that TgCPC1 is mainly localized in the PLVAC and the adjacent endosome-like compartment (ELC) through the use of two independent transgenic strains expressing endogenous epitope-tagged TgCPC1. Additionally, we generated a *TgCPC1*-null mutant in *Toxoplasma* parasites. Our data showed that the parasites use TgCPC1 to post-translationally modify some micronemal virulence factors, which are utilized for parasite invasion and egress. Collectively, our results elucidate a novel function of cathepsin C-like cystine exopeptidase within intracellular microbial pathogens for infections.

## RESULTS

### 1. TgCPC1 is localized within the endolysosomal system in *Toxoplasma* parasites

Cathepsin C protease, also named dipeptidyl peptidase I (DPP-I), is widely distributed throughout a variety of eukaryotes, including mammals and parasites. In mammalian cells, cathepsin C protease is located in the lysosome (24). *Toxoplasma* encodes three cathepsin C-like proteases in its genome (17). A previous report showed that TgCPC1 is localized within dense granules (17). However, a signal peptide sequence was predicted at the N-terminus of TgCPC1 (**Fig. S1A**), suggesting that the protease enters the parasite’s endolysosomal system. To determine the subcellular location of TgCPC1 in the parasites, we epitope-tagged TgCPC1 at two positions for immunofluorescence microscopy analysis (**Fig. 1B**). One strain contains a C-terminally 3xmyc-tagged TgCPC1, named TgCPC1-3xmyc^c^, while another strain expresses an internal 3xmyc tag, named TgCPC1-3xmyc^i^. The insertion site for the internal 3xmyc is preceded by a predicted antigenic region, as predicted by EMBOSS program, in order to ensure the epitope tag would be exposed on the surface of TgCPC1 for antibody detection by subsequent immunoblotting (IB) and immunofluorescence (IFA) assays (**Fig. S1B)**.

**Figure 1.**
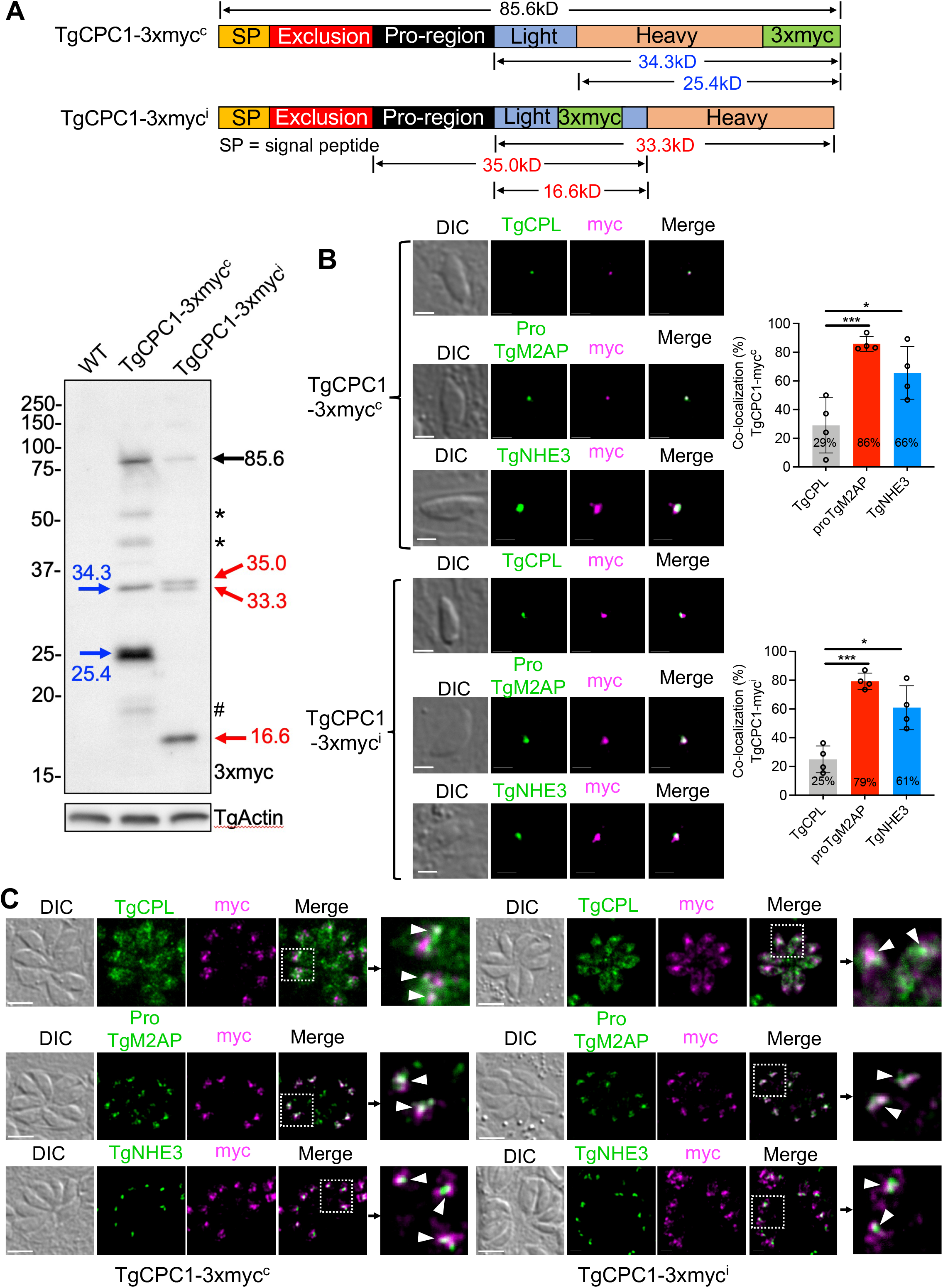
*Toxoplasma* cathepsin C-like protease 1 (TgCPC1) is an endolysosomal protease. (A) Schematic of epitope-tagging TgCPC1 expressed in *Toxoplasma* parasites. A 3xmyc tag was inserted at the C-terminus or within the light chain of TgCPC1 or at the C-terminus, which created TgCPC1-3xmyc^c^ and TgCPC1-3xmyc^i^ strains, respectively. Immunoblotting analysis showed that TgCPC1 is cleaved into a few species via multiple proteolytic cleavages. Based on the cleavage patterns of TgCPC1 seen in the immunoblots, TgCPC1 can be labeled into five domains. The domain division was deduced from the domain annotation of human cathepsin C protease via homologous alignment between TgCPC1 and human cathepsin C protease. The apparent molecular weights of TgCPC1 intermediates and final cleavage products were calculated based on their migration distances within SDS-PAGE. The intermediates and final products corresponding to individual molecular weights are annotated in the schematic. The polypeptides derived from TgCPC1-3xmyc^c^ and TgCPC1-3xmyc^i^ were marked in blue and red, respectively. The bands denoted by asterisks are degradation products. TgActin was probed as the loading control. (B) Both TgCPC1-3xmyc^c^ and TgCPC1-3xmyc^i^ strains were co-stained with antibodies recognizing the myc epitope and either the PLVAC marker (TgCPL) or ELC markers (TgNHE3 and proTgM2AP). Immunofluorescence microscopy (IFA) of pulse-invaded parasites revealed that ~70-75% of TgCPC1 is localized in the ELC, while ~25-30% of TgCPC1 resides within the PLVAC. Co-localization analysis was quantified in ~80 parasites per biological replicate for four independent trials. Bar = 2 µm. One-way ANOVA test was used to determine statistical significance; *, *p*<0.05; ***, *p*<0.001. (C) TgCPC1 was mainly located in the ELC within replicated parasites. Only a minute amount of TgCPC1 was observed to overlap with TgCPL. The co-localization between TgCPC1 with TgCPL (the PLVAC marker) or proTgM2AP/TgNHE3 (the ELC markers) were denoted by white arrowheads.

Protein lysates of both TgCPC1-3xmyc^c^ and TgCPC1-3xmyc^i^ strains were probed against anti-myc antibodies to confirm their expression. A few bands were observed by immunoblotting (**Fig. 1A**). In mammalian cells, cathepsin C is divided into five domains, signal peptide, exclusion region, propeptide region, heavy chain, and light chain (25). The light chain is followed by the heavy chain and both are cross-linked by disulfide bonds (25, 26). Similarly, multiple protein species of TgCPC1 were detected in both TgCPC1-3xmyc^c^ and TgCPC1-3xmyc^i^ parasites (**Fig. 1A**), indicating that TgCPC1 undergoes extensive processing within the parasites. The pro-form of TgCPC1, migrating at 85.6 kDa, was seen in both strains. The C-terminal 3xmyc-tagged TgCPC1-3xmyc^c^ strain displayed a major polypeptide chain migrating at 25.4 kDa, whereas the internally 3xmyc-tagged TgCPC1-3xmyc^i^ strain showed a smaller polypeptide migrating at 16.6 kDa as a predominant species. Additionally, TgCPC1-3xmyc^c^ has a single band of intermediate species at 34.3 kDa. In contrast, TgCPC1-3xmyc^i^ has a doublet band of intermediate species at 33.3 kDa and 35.0 kDa. The immunoblotting patterns suggest that the light chain of TgCPC1 precedes the heavy chain, given that the C-terminally tagged TgCPC1 species is larger than the internally 3xmyc-tagged band (**Fig. 1A and S1C**). This order is opposite to that is seen in the mammalian cathepsin C protease. Next, we performed IFA to determine the location of TgCPC1 within the parasite’s endolysosomal system by co-staining TgCPC1 with four endolysosomal markers, TgCPL (PLVAC marker), TgVP1 (PLVAC/ELC marker), proTgM2AP and TgNHE3 (ELC markers). In the pulse-invaded parasites, co-localization quantification between TgCPC1 and those markers indicated that the majority of TgCPC1 (~75%) is localized to the ELC, and the remaining 25% of TgCPC1 showed PLVAC localization (**Fig. 1B**). In replicated parasites, TgCPC1 is mainly localized to the ELC and only a minute amount of TgCPC1 appeared in the PLVAC (**Fig. 1C**). Since a previous report showed that TgCPC1 is localized in the dense granules (17), we stained the replicated TgCPC1-3xmyc^c^ and TgCPC1-3xmyc^i^ parasites with anti-TgGRA7 and anti-myc antibodies but did not observe staining of TgCPC1 within the PV (**Fig. S2**). We also tested the secretion of TgCPC1 by probing the constitutive excretory secreted antigen (ESA) fraction with antibodies recognizing the myc epitope, TgCPL (a PLVAC-localizing protein as a negative control), and TgPI-1 (a dense granule protein (27) as a positive control). Interestingly, we saw a very low level of secretion of TgCPC1 in ESA (**Fig. S3**). These findings suggest that a minute amount of TgCPC1 is routed to the default secretion pathway, although further investigation is needed to study the trafficking mechanism. Collectively, our data revealed that TgCPC1 is mainly located within the endolysosomal system in *Toxoplasma*.

### 2. TgCPC1 plays an important role in parasite invasion, egress, and migration

Given that TgCPC1 is located primarily within the ELC, we speculated that TgCPC1 is involved in cleaving other endolysosomal proteins in the parasites; therefore, the deletion of TgCPC1 would impair the parasite’s lytic cycle and virulence. To test this hypothesis, we genetically ablated the entire *TgCPC1* locus via homologous recombination to create a *TgCPC1*-null mutant, named Δ*cpc1* (**Fig. S4A**). To validate that the phenotypic defects observed in Δ*cpc1* are due to the loss of *TgCPC1*, a complementation plasmid containing the coding sequence of *TgCPC1* flanked by its 5’ and 3’ UTRs, as well as a bleomycin resistance cassette (**Fig. S4A**), was introduced into Δ*cpc1* to generate a Δ*cpc1CPC1* complementation strain. The TgCPC1 deletion and complementation were confirmed by PCR and quantitative PCR (qPCR) (**Fig. S4B and S4C**).

First, the general lytic cycle was assessed in WT, Δ*cpc1*, and Δ*cpc1CPC1* strains. The plaque number and area of Δ*cpc1* parasites are approximately 50% of that observed in WT and Δ*cpc1CPC1* parasites (**Fig. 2A**). Interestingly, the plaques derived from Δ*cpc1* were filled with lysed parasites (**Fig. 2A**), suggesting that the *TgCPC1*-deletion mutant cannot migrate as efficiently as WT parasites, further affecting its lytic cycle. To test this, we quantified the form and velocity of parasite movement via live imaging and found that the percentage of circular motility of Δ*cpc1* was reduced by 50% compared to WT and Δ*cpc1CPC1* parasites (**Fig. 2B**). Similarly, the velocity of the movement of Δ*cpc1* parasites was ~50% of that seen in WT and Δ*cpc1CPC1* (**Fig. 2B**). These findings led to our next assessment of parasite invasion and egress, which are steps requiring efficient parasite movement. To compare invasion efficiency in these strains, the parasites were overlayed onto confluent human foreskin fibroblasts (HFFs) for 30 min prior to immunostaining of extracellular and intracellular parasites. Our results showed that the invasion of Δ*cpc1* parasites was decreased by 50% relative to WT and Δ*cpc1CPC1* (**Fig. 2C**). To quantify egress in these strains, infected HFFs were stimulated by zaprinast for 5 min to induce parasite egress, which results in the release of lactate dehydrogenase (LDH) from host cells. The quantity of LDH released is proportional to parasite egress efficiency. Our results showed that the Δ*cpc1* parasites decreased egress by ~50% compared to WT and Δ*cpc1CPC1* parasites (**Fig. 2D**). Small plaques can also be a result of reduced replication. To test this, we grew WT, Δ*cpc1*, and Δ*cpc1CPC1* parasites in HFFs and quantified the number of parasites per individual PVs. At 28 hrs post-infection, the Δ*cpc1* strain did not show any growth defects (**Fig. 2E**). To investigate whether these defects throughout the lytic cycle in Δ*cpc1* lead to reduced acute virulence, outbred CD-1 mice were injected subcutaneously with 100 WT, Δ*cpc1*, or s Δ*cpc1CPC1* parasites and monitored daily for symptoms over the course of a 30-day period. The mice infected with Δ*cpc1* survived significantly longer than mice infected with WT or Δ*cpc1CPC1* parasites via subcutaneous injection (**Fig. 2F**). Interestingly, one mouse infected with Δ*cpc1CPC1* survived for 18 days post-infection, but this survival difference between WT and Δ*cpc1CPC1* was not statistically significant (**Fig. 2F**). Taken together, these results indicate an important role of TgCPC1 in parasite invasion, egress, and migration, but not replication. Furthermore, TgCPC1 is required for optimal infection of *Toxoplasma* parasites.

**Figure 2.**
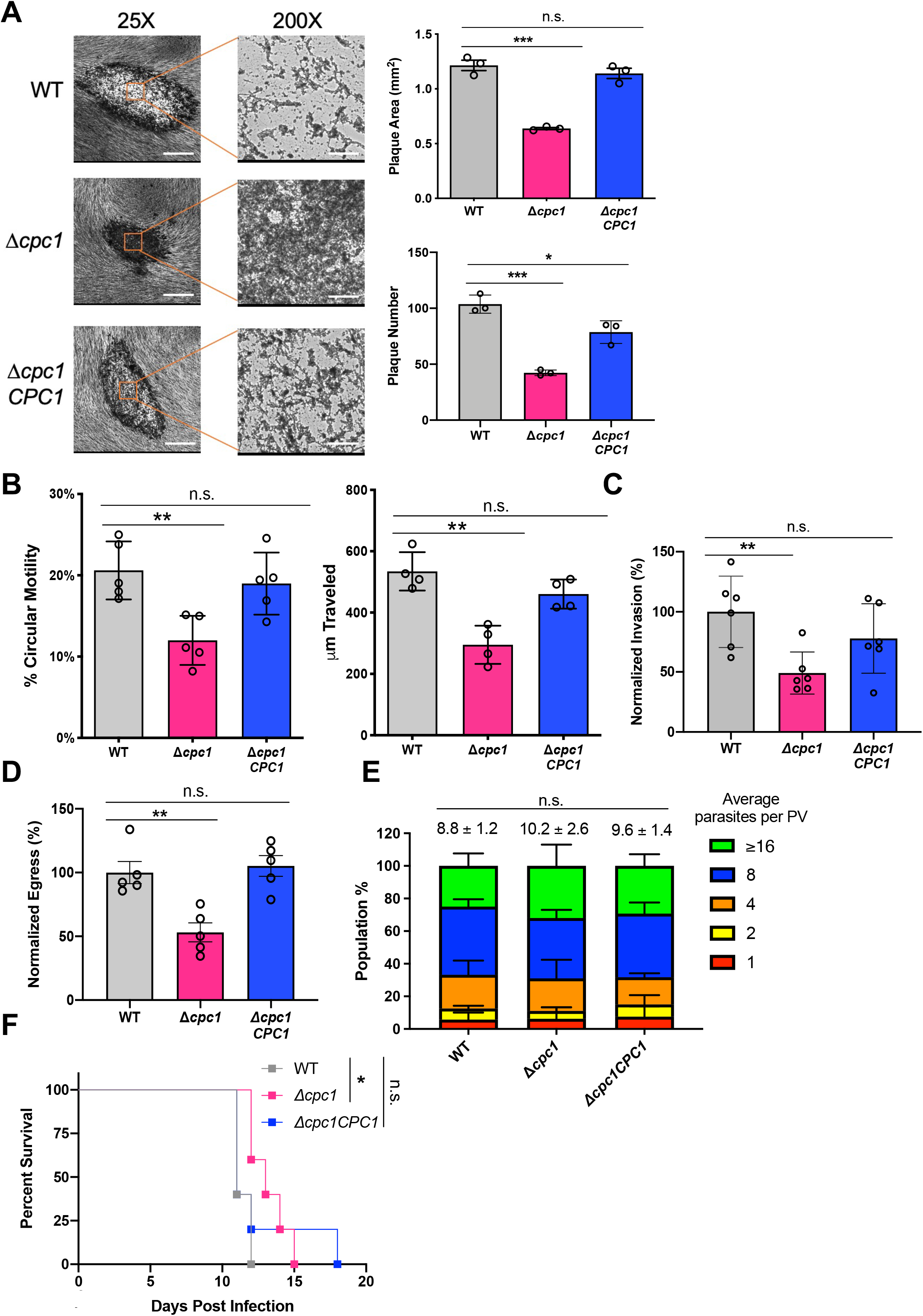
TgCPC1 plays an important role in the lytic cycle of *Toxoplasma* parasites and their acute virulence. (A) The *TgCPC1*-deletion mutant displayed fewer and smaller plaques than WT and Δ*cpc1CPC1* parasites. A noteworthy characteristic of Δ*cpc1* plaques is the lack of a clear central region, suggesting that the mutant cannot migrate efficiently. Three independent assays were completed. Statistical analysis was completed using one-way ANOVA and WT was used as the control for comparison. Bar = 500 µm and 50 µm in the 25x and 200x amplification images, respectively. (B) Parasite motility was chemically induced by adding 100 mM zaprinast and recorded by time lapse videos using an inverted fluorescence microscope with a CCD camera. The circular motility and the total distance traveled revealed that the motility of the Δ*cpc1* parasites was significantly reduced compared to WT and Δ*cpc1CPC1*. Data shown here were derived from at least four independent trials. One-way ANOVA was used for statistical analysis. (C) Parasite invasion was reduced by ~50% in Δ*cpc1* compared to WT and Δ*cpc1CPC1*. Six fields of view were counted for each strain per biological replicate in a total of six individual trials. (D) Lactate dehydrogenase release-based egress assay revealed that egress in Δ*cpc1* was reduced by ~50% compared to WT and Δ*cpc1CPC1*. Data from five trials were combined for statistical calculation. (E) Replication assays were performed by quantifying the number of parasites per PV in WT, Δ*cpc1*, and Δ*cpc1CPC1* at 28 hrs post-infection. One hundred PVs were enumerated per replicate in a total of three replicates and plotted. The average numbers of parasites for individual strains were compared for statistical significance calculation. All strains displayed comparable replication rates. Statistical significance for assays listed in panels C through E were determined using unpaired Student’s *t*-test. (F) Acute virulence was evaluated in a murine model via subcutaneous infection. One hundred parasites from each strain were used to infect outbred CD-1 mice (n=5 per strain). Mice infected with Δ*cpc1* had a modest yet significant increase in survival time. Data were recorded and presented using the Kaplan-Meier plot. Statistical analysis was performed using the Log-rank (Mantel-Cox) test. *, *p*<0.05; **, *p*<0.01; ***, *p*<0.001; n.s., not significant.

### 3. Parasites lacking *TgCPC1* displayed altered protein secretion

During the lytic cycle, the parasites secrete micronemal effectors to facilitate parasite invasion and egress. For example, TgMIC2 and TgM2AP are involved in parasite invasion (28–30) while TgPLP1, a perforin-like protein, is released by parasites for egress (31). Given the invasion, egress, and migration defects observed in Δ*cpc1*, we assessed if the mutant parasites showed abnormal microneme secretion. To test this, we liberated WT, Δ*cpc1*, and Δ*cpc1CPC1* from host cells and prepared ESA fractions to quantify microneme secretion. In both constitutive and induced ESA fractions, the migration patterns of several microneme proteins were altered. In WT parasites, TgMIC2 showed two species in the ESA migrating at 95 kDa and 100 kDa, whereas only one TgMIC2 species at 100 kDa was observed in Δ*cpc1* (**Fig. 3A**). Similarly, TgM2AP underwent a few proteolytic modifications on the surface in WT parasites, which is subsequently secreted into the ESA fraction. However, the majority of secreted TgM2AP in Δ*cpc1* was accumulated as pro- and mature forms of TgM2AP and a series of smaller cleaved TgM2AP species were not observed (**Fig. 3A**). The mature form of TgM2AP in Δ*cpc1* is slightly bigger than that in WT parasites. Similar to TgM2AP, the mature form of TgAMA1 in Δ*cpc1*, another key invasion micronemal effector (32, 33), migrated slowly relative to that in WT parasites (**Fig. 3A**), suggesting that TgCPC1 is involved in the processing of the full-length TgAMA1 into its pro-form. The secreted ecto TgAMA1 in Δ*cpc1* was also bigger than that in WT and Δ*cpc1CPC1* parasites (**Fig. 3A**), suggesting that TgCPC1 mediates the formation of mature TgAMA1 before they are cleaved by TgROM4 within the plasma membrane. The migration pattern of TgMIC5 in both WT and Δ*cpc1* was similar; however, we observed increased secretion in Δ*cpc1* parasites (**Fig. 3A**). TgMIC5 remained in the unprocessed pro-form to a greater extent in Δ*cpc1* in the constitutive ESAs compared to WT and Δ*cpc1CPC1*, but this was not observed in the induced ESAs (**Fig. 3A**). As a main egress effector, TgPLP1 is proteolytically processed into a few smaller species whose molecular weights migrate around 95 kDa (5, 34). We found that the abundance of these proteolytically processed species of TgPLP1 were significantly decreased in Δ*cpc1* and instead, TgPLP1 accumulated predominantly as a polypeptide migrating at approximately 130 kDa (**Fig. 3A**). Interestingly, we also detected higher secretion of dense granules in the Δ*cpc1* parasites, such as TgGRA7 and TgPI-1 (protease inhibitor-1) proteins (**Fig. 3B**). Hence, the deletion of *TgCPC1* globally changes protein secretion in *Toxoplasma* and alters the migration patterns of several critical invasion and egress effectors, such as TgMIC2, TgM2AP, TgAMA1, and TgPLP1.

**Figure 3.**
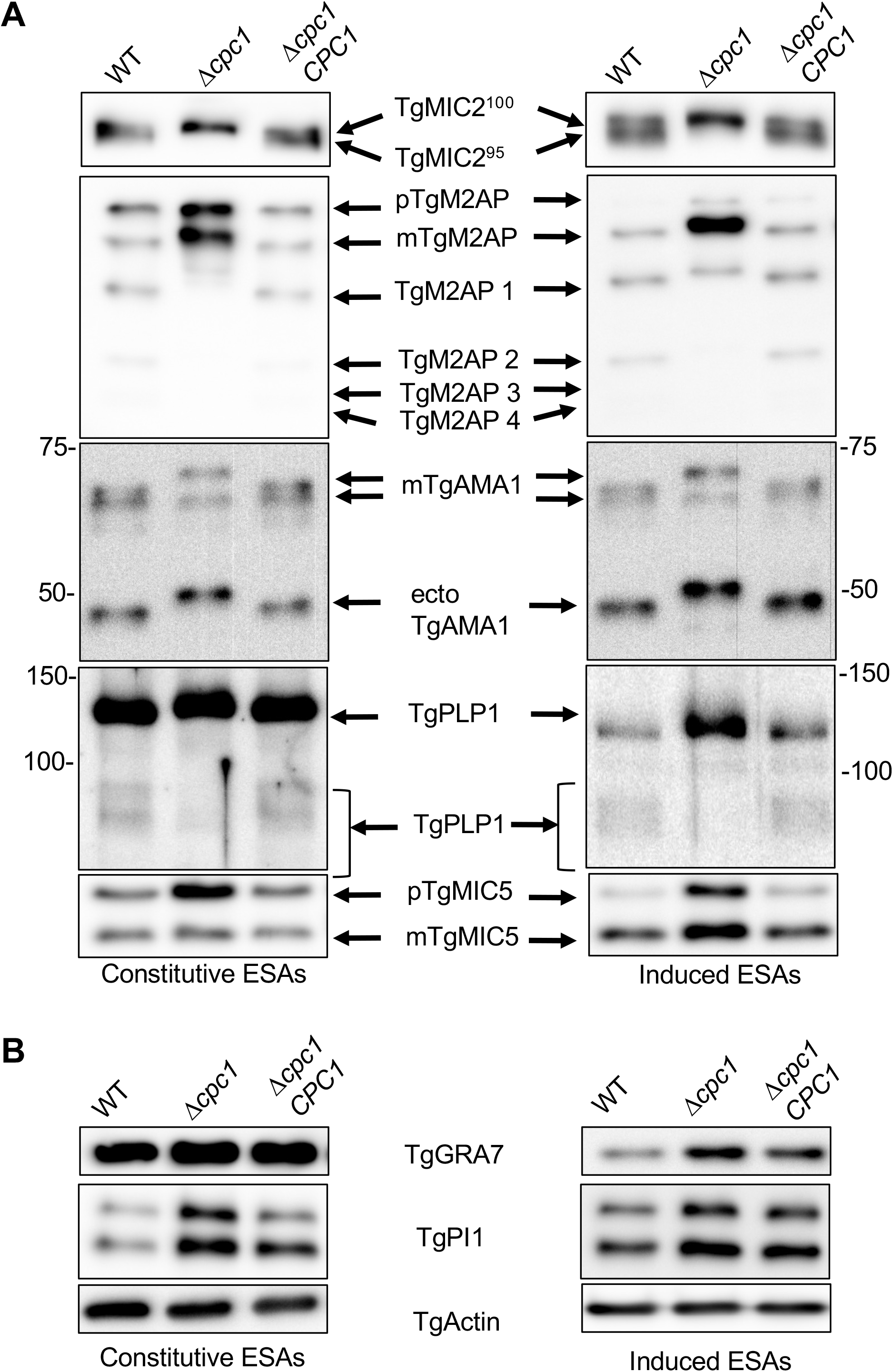
The protein secretion patterns were altered in *Δcpc1*. (A) Several microneme proteins were not properly trimmed and released in excretory secretory antigen (ESA). ESA fractions were prepared by standard constitutive and 1% ethanol-induced protein secretion. Purified ESAs were probed against a few representative microneme proteins, such as TgMIC2, TgM2AP, TgAMA1, TgPLP1, and TgMIC5. (B) Evaluation of dense granule secretion in *TgCPC1*-deficient parasites via immunoblotting. TgActin was probed against the lysates as loading controls. At least three independent preparations of constitutive and induced ESA samples were generated for this assay.

### 4. Defective microneme secretion in Δ*cpc1* was not caused by abnormal protein trafficking or altered total protein abundance

The migration patterns of several microneme proteins, which traffic through the endolysosomal system before arriving at microneme, were altered within Δ*cpc1* parasites. The lack of TgCPC1 may alter the subcellular trafficking or affect the total abundances of these proteins, which will ultimately affect their downstream secretion.

First, we probed the lysates from WT, Δ*cpc1*, and Δ*cpc1CPC1* parasites to assess the total abundance of micronemal proteins via immunoblotting. The protein levels of TgMIC2 in all lysates were comparable (**Fig. 4A**). In WT parasites, TgM2AP exists as two forms, pro- and mature (pTgM2AP and mTgM2AP, respectively), which are produced by proteolysis during intracellular trafficking (35, 36). The total level of TgM2AP was slightly increased in the Δ*cpc1* parasites. More strikingly, the mTgM2AP species in Δ*cpc1* migrated slightly slower than that in WT and Δ*cpc1CPC1* (**Fig. 4A**). Similarly, we also observed that the mature TgAMA1 in Δ*cpc1* was slightly larger than that in WT and Δ*cpc1CPC1* strains (**Fig. 4A**). Previous work has revealed both TgASP3 and TgCPL are involved in the conversion of pTgM2AP into mTgM2AP (1, 2), but it still remains unknown about the protease(s) for TgAMA1 maturation. Our data suggest that TgCPC1 is involved in the final maturation of TgM2AP and TgAMA1. For TgPLP1, the species observed at ~130 kDa also migrated slowly in the Δ*cpc1* parasites. In addition, we also observed a few cleaved TgPLP1 bands migrating slowly in the lysates of Δ*cpc1*, compared to WT and Δ*cpc1CPC1* (**Fig. 4A**). Interestingly, we observed an enhanced abundance of TgMIC5 levels in the Δ*cpc1* lysate. Like TgM2AP, TgMIC5 underwent proteolytic cleavage for the formation of mTgMIC5. In Δ*cpc1*, the ratio of the pro-form of TgMIC5 over the mature form is decreased (**Fig. 4A**).

**Figure 4.**
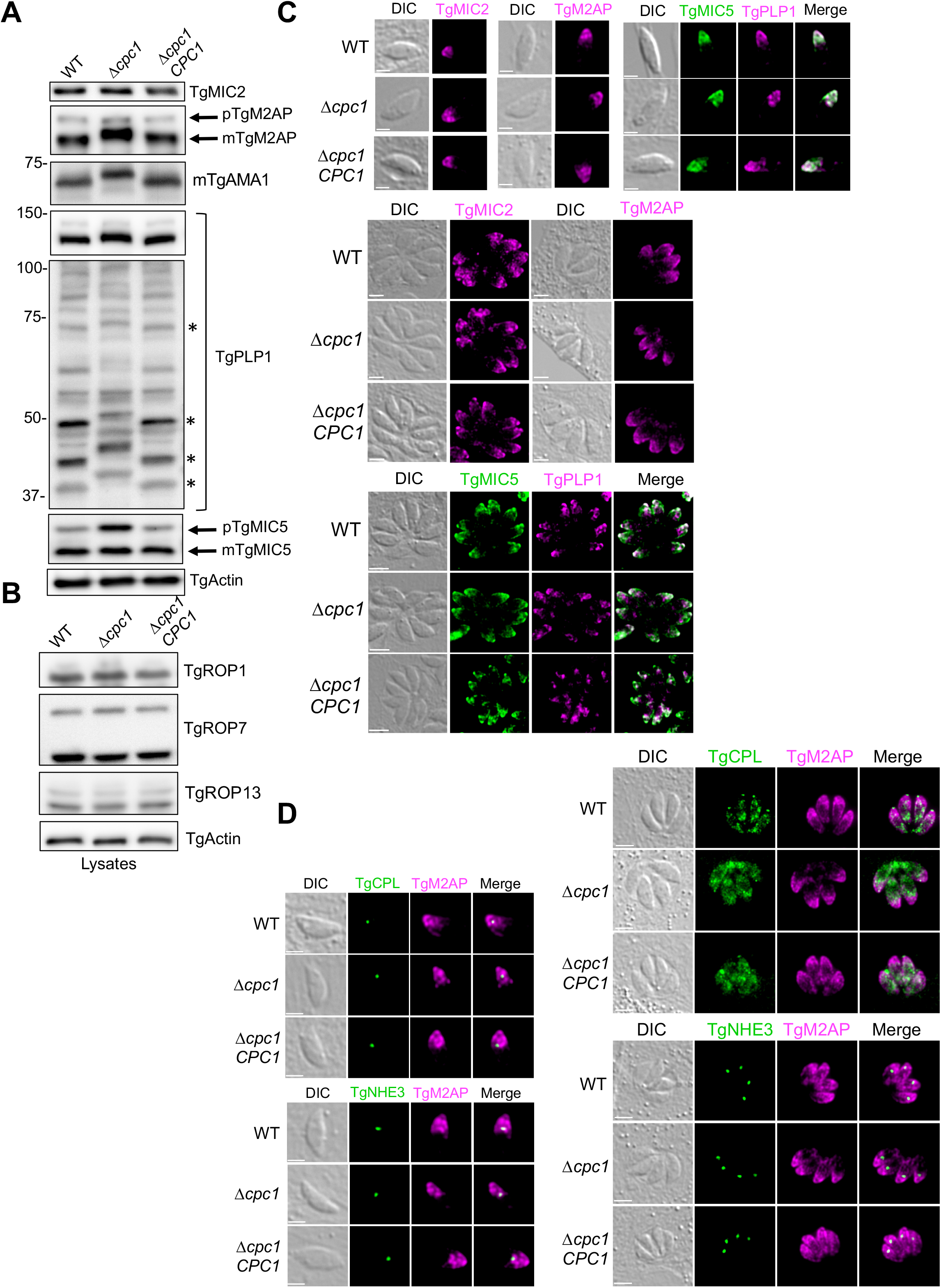
Intracellular trimming of some micronemal proteins was altered in Δ*cpc1*, while their intracellular trafficking was not changed. (A) The micronemal proteins probed in Fig. 3A were also probed in the lysates to assess the roles of TgCPC1 in micronemal protein trimming. (B) A few rhoptry proteins were also probed in the lysates to assess if TgCPC1 is involved in rhoptry protein maturation. (C) Some representative micronemal proteins were stained in pulse-invaded and replicated WT, Δ*cpc1*, and Δ*cpc1CPC1* parasites to test if defective intracellular trimming impairs their delivery to the micronemes. (D) The lack of TgCPC1 cleavage in microneme protein trimming did not lead to abnormal accumulation in the ELC and PLVAC. Bar = 2 µm or 5 µm in pulse-invaded and newly replicated parasites, respectively in (C) and (D). All assays were repeated at least in triplicate.

Rhoptry proteins also traffic through the ELC (1, 37, 38). To test if TgCPC1 is involved in modifying rhoptry proteins within the endolysosomal system, we also evaluated the total abundance and proteolytic processing patterns of TgROP1, TgROP7, and TgROP13 by immunoblotting. We did not see any changes in these representative rhoptry proteins (**Fig. 4B**). To further test if the incompletely trimmed microneme proteins undergo normal subcellular trafficking, we immunostained, pulse-invaded, and replicated parasites with TgMIC2, TgM2AP, TgMIC5, and TgPLP1 antibodies. We observed typical microneme staining for these proteins, located at the apical end of the parasites (**Fig. 4C**), indicating that the final trimming of microneme proteins is not essential for their delivery to micronemes. Further, we co-stained, pulse-invaded, and replicated parasites with anti-TgM2AP antibody, as well as serum recognizing PLVAC (anti-TgCPL) and ELC (anti-TgNHE3) markers to determine if the incompletely maturated microneme proteins accumulated within the endolysosomal system. As expected, our IFA data showed that some of TgM2AP proteins co-localized with TgNHE3 within the ELC but not with TgCPL. This observation is consistent with a previous report, showing that the ELC is a site for the maturation of some microneme proteins by TgCPL (2). However, we did not observe a greater accumulation of TgM2AP in the ELC in Δ*cpc1* parasites compared to WT parasites (**Fig. 4D**), suggesting that the additional residues at the N-terminal end of mTgM2AP that are cleaved by TgCPC1 do not impair its subcellular trafficking. Taken together, these results indicate that in the absence of TgCPC1, some microneme proteins cannot be fully processed despite being able to properly traffic to micronemes.

### 5. Defective microneme protein secretion in Δ*cpc1* is due to the blocked maturation of TgSUB1

TgSUB1, a subtilisin-like micronemal protease, traffics to and remains at the parasite’s cell surface via a GPI-anchor (5, 39). It has been reported that TgSUB1 plays a major role in the processing of micronemal effectors for ESA secretion (5). Given that TgMIC2 in the Δ*cpc1* ESA was only observed in the TgMIC2^100^ form and a series of processed TgM2AP species were lost, mirroring the phenotype observed in Δ*sub1* (5), we speculated that TgSUB1 is not maturated correctly within Δ*cpc1*. To test this hypothesis, we prepared constitutive and induced ESAs and probed them against anti-TgSUB1 antibody for immunoblotting. As expected, TgSUB1 in Δ*cpc1* migrated as the ~90 kDa pro-form version, while the majority of TgSUB1 protein in WT and Δ*cpc1CPC1* migrated at ~70 and 82 kDa (**Fig. 5A**). A similar observation was seen in the induced ESAs (**Fig. 5A**). Furthermore, we probed the parasite lysates against anti-TgSUB1 and found that that the TgSUB1 protein cannot be cleaved within Δ*cpc1* (**Fig. 5A**), suggesting that TgCPC1 plays an essential role in the maturation of TgSUB1. To test if the pro-form of TgSUB1 could be delivered to the surface of Δ*cpc1* parasites, we immunostained filter-purified extracellular parasites with anti-TgSUB1 antibody, in the absence of cell membrane permeabilization by Triton X-100, and saw comparable staining of surface-localized TgSUB1 in Δ*cpc1* relative to WT (**Fig. 5B**). To further evaluate the subcellular trafficking of the pro-form of TgSUB1 in the parasites, pulse-invaded and replicated WT and Δ*cpc1* parasites were subjected to IFA analysis by co-immunostaining with antibodies recognizing TgSUB1 and TgMIC5. TgMIC5 serves as a microneme marker since we previously showed its maturation pattern was unchanged in Δ*cpc1*. Staining of both TgSUB1 and TgMIC5 was well co-localized within the Δ*cpc1* parasites (**Fig. 5C**), indicating that the inability of maturating TgSUB1 did not impair its delivery to micronemes. A previous report found that when the propeptide of TgSUB1 was fused at the N-terminal end of GFP, the chimeric protein was retained to the ELC (40). Therefore, we speculated that the incorrectly trimmed TgSUB1 in Δ*cpc1* may be retained in the ELC to a greater extent than that in WT. To test this, we assessed the extent to which TgSUB1 co-localized within the PLVAC or ELC in the parasites by using TgCPL and TgNHE3 as PLVAC and ELC markers, respectively. Some TgSUB1 staining co-localized with TgNHE3 in the ELC within both pulse-invaded and replicated WT, Δ*cpc1*, and Δ*cpc1CPC1* parasites to a similar extent, while TgCPL fragmented during intracellular replication and did not co-localize with TgSUB1 (**Fig. 5D**). These results revealed that the presence of TgCPC1 is essential for the maturation of TgSUB1 protease, but full blockage of TgSUB1 maturation does not alter its intracellular trafficking and distribution on the parasite’s cell surface. However, the inactive form of TgSUB1 trafficked to the parasite’s surface is unable to carry out the surface processing of other microneme proteins such as TgMIC2, TgM2AP, and TgPLP1, which leads to defects in parasite invasion and egress.

**Figure 5.**
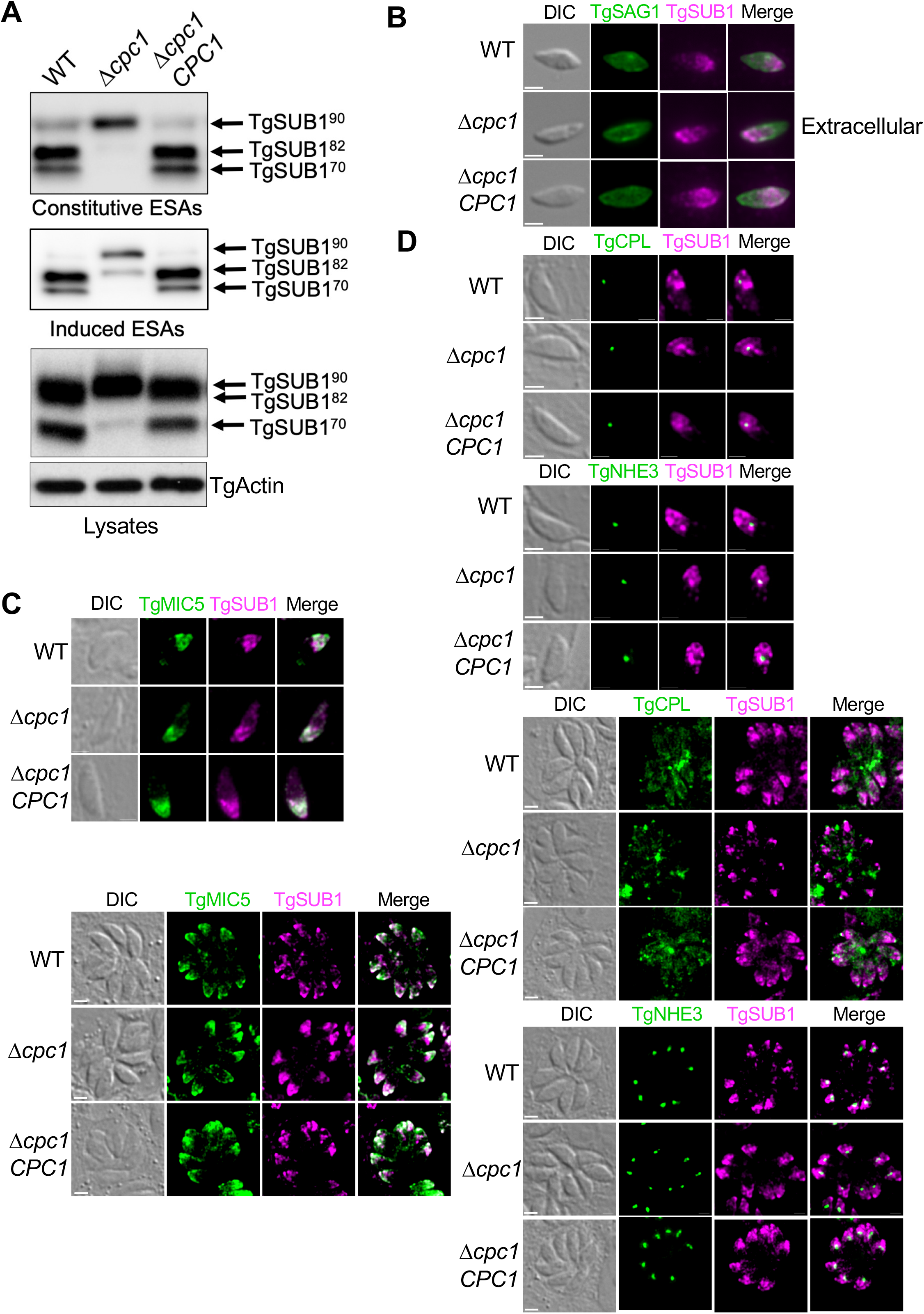
Altered microneme protein secretion in Δ*cpc1* is due to blocked maturation of TgSUB1. (A) Constitutive and induced ESAs as well as lysates from WT, Δ*cpc1*, and Δ*cpc1CPC1* were probed with a TgSUB1-recognizing antibody. TgSUB1 cannot be maturated into its mature form in Δ*cpc1* parasites. Accordingly, the TgSUB1 on the parasite surface is not active within the *TgCPC1*-deletion mutant. (B) To evaluate the abundance of surface-localized TgSUB1, extracellular, non-permeabilized parasites were immunostained and imaged. TgSAG1 was included as a positive control. Immunofluorescence microscopy revealed that TgSUB1 still trafficked normally to the surface of Δ*cpc1* parasites, albeit in an inactive form. (C) TgSUB1 staining in fully permeabilized, pulse-invaded and replicated Δ*cpc1* mutant parasites showed that the immature TgSUB1 still traffics to the micronemes properly. Bar = 2 µm. (D) Some TgSUB1 accumulated in the ELC prior to trafficking to micronemes. The loss of TgCPC1 blocked the maturation of TgSUB1 but did not result in its accumulation in the ELC to a greater extent than that in WT and Δ*cpc1CPC1*. Bar = 2 µm.

### 6. Chemical inhibition of TgCPC1 recapitulated phenotypes seen within Δ*cpc1*

A BLAST search revealed TgCPC1 as an ortholog of the *Plasmodium falciparum* dipeptidyl aminopeptidase (PfDPAP1; PF3D7_1116700). PfDPAP1 has been reported as an attractive drug target (41). A few small chemical inhibitors have been shown to have high potencies against PfDPAP1 (42). A recent initiative “opnMe” (www.opnMe.com) for sharing resources used in biomedical research reported that a chemical inhibitor, named BI-2051, is a selective, soluble, and cell-permeable inhibitor for PfDPAP1 with an IC_50_of 0.3 nM. Its inhibition against human cathepsin C protease, termed dipeptidyl aminopeptidase I (hDPP-I), is ~10-fold less than that observed in recombinant PfDPAP1 (opnMe). To test if TgCPC1 is targeted by BI-2051, infected HFFs were incubated with 10, 1, or 0.1 μM BI-2051 or with the DMSO vehicle control, for a plaque assay. Only at 10 µM BI-2051, the plaques formed by WT parasites were significantly smaller than un-treated samples (**Fig. 6A**). But the number of plaques formed in Δ*cpc1*-infected host cells was comparable to that in WT (**Fig. 6A**), suggesting that the inhibition of TgCPC1 by BI-2051 will not take effect in a short timeframe *in vivo*. Similar to the Δ*cpc1* plaques, the BI-2051-treated WT plaques were filled with lysed parasites (**Fig. 6A**), suggesting that the treated parasites have reduced motility. To test if this cathepsin C inhibitor could block intracellular TgSUB1 maturation and further microneme processing on the parasite’s surface, WT *Toxoplasma* parasites were grown in HFFs in the presence of 10 µM BI-2051 for 48 hrs before filter-purification and lysate preparation. The DMSO-treated WT and Δ*cpc1* were included as negative and positive controls, respectively. Similar to the phenotypes observed in Δ*cpc1*, the maturation of TgSUB1 was significantly blocked and the mature form of TgM2AP migrated slightly slowly relative to that shown in WT (**Fig. 6B**). Accordingly, most of the secreted TgSUB1 was retained as the immature form and the formation of TgM2AP1 that is processed by TgSUB1 was dramatically reduced **(Fig. 6B**). Collectively, chemical interrogation of TgCPC1 recapitulated the phenotypic defects observed in Δ*cpc1*, indicating that the BI-2051 inhibits TgCPC1 activity, albeit with reduced potency.

**Figure 6.**
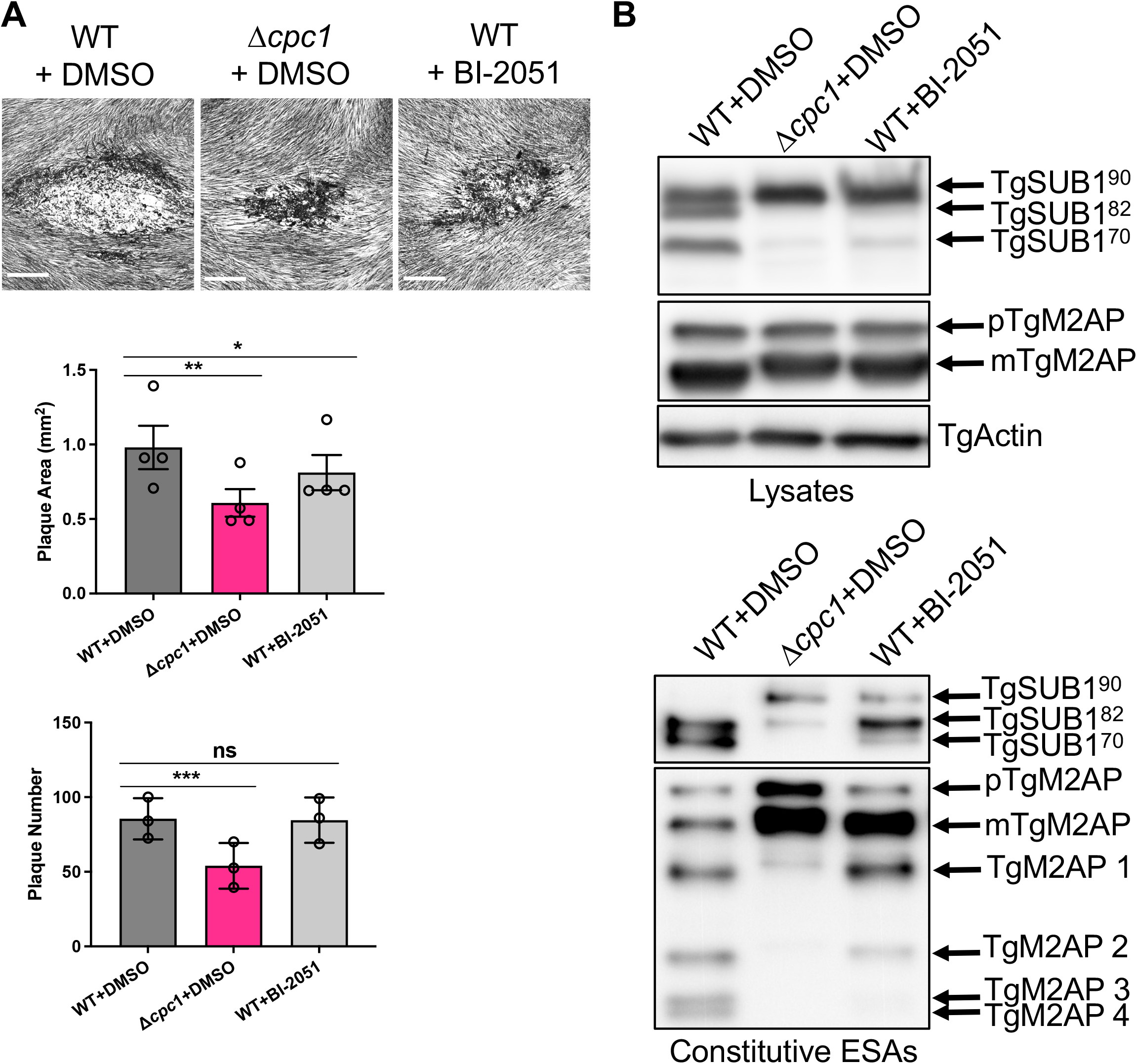
Chemical inhibition of TgCPC1 recapitulated the phenotypes seen within *Δcpc1*. WT parasites were treated with 10 µM BI-2051, a potent inhibitor against PfDPAP1, for 48 hrs before (A) plaque assay and (B) the preparation of lysates and ESAs. The plaque assay and immunoblotting showed that the proteolytic activity of TgCPC1 is important for the parasite’s lytic cycle, TgSUB1 maturation, and the final trimming of TgM2AP. Bar = 500 µm. Statistical significance in panel A was calculated by unpaired Student’s *t*-test. *, *p*<0.05; **, *p*<0.01; ***, *p*<0.001; n.s., not significant.

In contrast to its high potency against the malarial DPAP1 enzyme, BI-2051 only showed moderate potency targeting the *Toxoplasma* ortholog. To help elucidate the molecular mechanism by which BI-2051 interacts with PfDPAP1 and TgCPC1 proteins at the atomic level, we used Autodock Vina to perform a molecular docking simulation between the ligand and the structures of PfDPAP1 and TgCPC1 proteins predicted by the AlphaFold algorithm (43). We first docked BI-2051 at the active site of human cathepsin C protease, termed dipeptidyl aminopeptidase I (hDPP-I), since it was used as a model protein for aligning the coordinates of the active site residues of PfDPAP1 and TgCPC1 (**Fig. 7A**). As shown in **Figure 7B**, BI-2051 binds to the active site of hDPP-I with a binding affinity of–6.8 kcal/mol. As seen in the hDPP-I/BI-2051 binding pose, BI-2051 interacts with the amino acid residues; Asp-1, Gln-228, Cys-234, Gly-277, and Asn-380, similar to Gly-Phe-diazomethane co-crystallized with hDPP-I (26). These amino acids are conserved among all three orthologs and take part in the catalytic mechanism or substrate binding (**Fig. S5**) (26). The docking models showed that BI-2051 displays similar binding interactions with the conserved amino acids in all three cathepsin C orthologs. BI-2051 strongly binds to the active site of PfDPAP1 with a strong binding energy at–8.7 kcal/mol, whereas its binding affinity with TgCPC1 is dampened to–7.4 kcal/mol (**Fig. 7B**), although it is significantly lower than that of the interaction between BI-2051 with hDPP-I. This docking result is consistent with the experimental assays, which reveal BI-2051 as a more potent inhibitor against PfDPAP1 than TgCPC1 and hDPP-I. These findings suggest that there are structural differences between TgCPC1 and PfDPAP1 within these two representative apicomplexan parasites, indicating a potential for the development of specific inhibitors targeting cathepsin C protease which can be used for controlling apicomplexan parasite infections.

**Figure 7.**
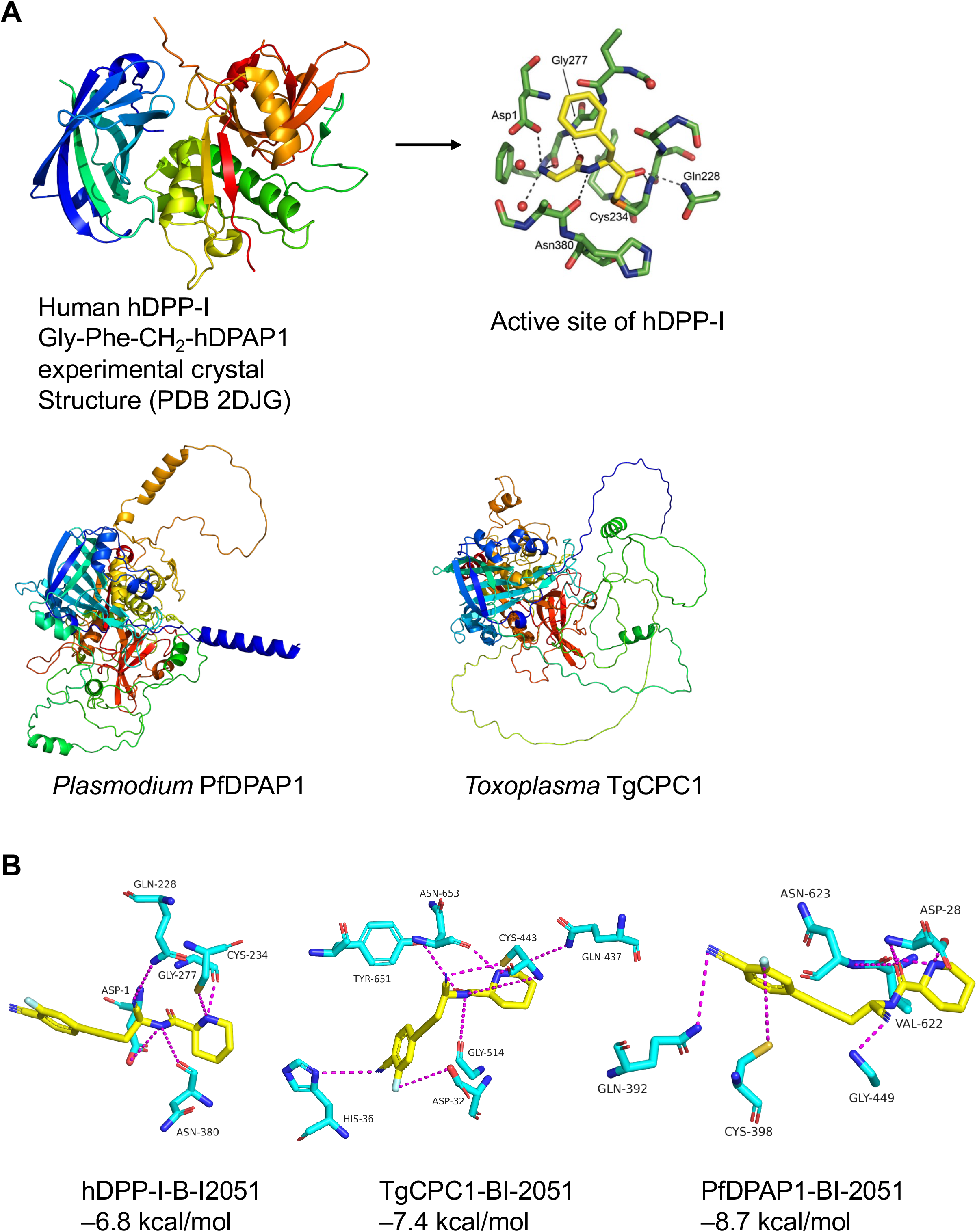
Molecular modeling of cathepsin C proteases with BI-2051. (A) The three-dimensional structure of human cathepsin C protease (hDPP-I) was acquired from the RCSB Protein Data Bank (PDB). The primary citation of related structures is 2DJG. The structures of *Plasmodium* and *Toxoplasma* orthologs were generated by Alpha-Fold algorithm. The coordinates of the active site residues of PfDPAP1 and TgCPC1 were predicted by sequence alignment with hDPP-I. (B) The BI-2051 was docked to the active sites of the proteins using AutoDock vina. The binding affinity of the ligand was reported in kcal/mol.

## DISCUSSION

Mining of the *Toxoplasma* genome reveals that there are five cysteine cathepsin proteases. Two of them, cathepsin L- and B-like proteases (TgCPL and TgCPB, respectively), have been located in the PLVAC (11). Three cathepsin C-like proteases were reported to be expressed at different infection stages (17). In contrast to their mammalian ortholog that resides in the lysosome, two independent 3xmyc-tagged TgCPC1 strains engineered in this study showed that most of TgCPC1 is located in the ELC, a precursor organelle to the PLVAC. TgCPC1 was previously reported to be a dense granule protease by IFA (17). Interestingly, the malarial ortholog of TgCPC1, named PfDPAP1, was reported as a food vacuole-residing protease and has also been seen in the PV (20). We co-immunostained replicated parasites using anti-myc epitope antibody along with anti-TgGRA7 serum and did not observe TgCPC1 staining in the PV (**Fig. S2**). However, we detected trace amounts of TgCPC1 secretion in the ESA (**Fig. S3**), suggesting that a minute amount of TgCPC1 may be secreted to the PV, albeit below the detection limits of IFA (**Fig. S2**). Within previous literature, it has been hypothesized that TgCPC1 protease can digest proteins in the PV for the parasite’s nutritional needs. which was assessed by treating replicated parasites with the cathepsin C inhibitor (17). The growth assay in our study did not show growth defects in the *TgCPC1*-deficient parasites, undermining this possibility. However, it remains possible that the PV-localizing TgCPC1 is involved in the process of egress by an underdetermined mechanism. In addition, a small amount of TgCPC1 may also be further released into host cells, in the same manner as other dense granule proteins, to modulate the host’s response. Previous work showed that the exogenous expression of TgCPC1 in HEK293 cells inhibits the NFΚB signaling (44), although it remains unknown if TgCPC1 can cross the parasitophorous vacuole membrane (PVM) into the host. Some microbial pathogens secrete proteases to assist in their invasion of host cells. For example, *Streptococcus pyogenes* releases SpeB, a cysteine protease, to degrade the host defense system, such as extracellular matrix and immune effectors (45). *Entamoeba histolytica* also uses cathepsin-like proteases to disrupt host cells for its infections (46). Therefore, the small amount of secreted TgCPC1 protein may aid in invasion and possibly in egress, as well. More evidence is needed to support this speculation. Interestingly, a recent hyperLOPIT (localization of organelle proteins by isotope tagging) proteomic analysis for *Toxoplasma* proteins revealed that the TgCPC1 protein was grouped with microneme proteins (23), which supports our finding of its secretion within ESAs. However, our IFA assay did not reveal TgCPC1 within micronemes, probably due to its extremely low abundance.

The mature form of mammalian cathepsin C protease is composed of two chains linked by disulfide bonds (25). The heavy chain, containing the active cysteine residue, is preceded by the light chain, where the essential histidine and asparagine are located. The protein sequence alignment between *Toxoplasma* and human orthologs revealed the potential cleavage sites (**Fig. S1C**) within the primary sequence of *Toxoplasma*. Based on the migration patterns of the cleaved proteins, our immunoblotting results suggested that *Toxoplasma* CPC1 exhibits an opposite arrangement of the heavy and light subunits (**Fig. 1A**). Given the locations of the 3xmyc epitopes that were engineered within the primary structure of TgCPC1, the active cysteine residue is located in the putative light chain based on our immunoblotting results (**Fig. S1C**). It is unclear why *Toxoplasma* adopts a different strategy for the arrangement of both subunits within the mature TgCPC1 enzyme in comparison to the mammalian ortholog structure.

In mammalian cells, cathepsin L and S are potentially involved in cathepsin activation (47). However, the activation was still observed in the cathepsin L- and S-deletion cell lines, suggesting that other protease(s) participate in the proteolytic processing (47). To understand the relationship between cathepsin L and cathepsin C in *Toxoplasma*, we compared the processing patterns of TgCPC1 in WT and Δ*cpl* parasites. Interestingly, we did not observe any alterations in the migration of TgCPC1-related bands in Δ*cpl*. Instead, the total amount of TgCPC1 was increased significantly in Δ*cpl*, regardless of the locations of the epitope tags (**Fig. S6A**). A similar phenotype was mirrored within WT parasites treated with LHVS, an inhibitor against TgCPL in *Toxoplasma* (**Fig. S6B**) (48). These findings suggest that TgCPL is involved in the homeostasis of TgCPC1, but other proteases probably mediate the cleavage of TgCPC1. The maturation location of TgCPC1 within the parasites remains to be determined.

The mutant parasites lacking TgCPC1 showed reduced invasion and egress but replication was unaffected. The loss of TgCPC1 resulted in significantly altered patterns of many micronemal proteins in the ESA fractions, including TgMIC2 and TgPLP1, two important virulence factors for parasite invasion and egress, respectively, due to the maturation of TgSUB1 being completely abolished. The loss of TgCPC1 also altered the maturation pattern of TgM2AP and TgAMA1, which suggests that TgCPC1, an aminopeptidase, conducts the final trimming of these micronemal effectors. Another interesting observation is that the ratio of the pro-form of TgMIC5 to its mature form was altered, indicating its proteolytic maturation is also impacted by TgCPC1. In addition, more TgMIC5 was secreted into the ESAs in Δ*cpc1*. Our previous work revealed that TgMIC5 mimics the pro-domain of TgSUB1 to regulate its proteolytic activity (49). In the Δ*cpc1* parasites, the pro-peptide of immature TgSUB1 is still associated with its mature form, which will block binding of TgMIC5 to the parasite’s surface, thus leading to increased secretion in the ESAs as we observed. The abnormal maturation of TgSUB1 may also affect the maturation patten of TgMIC5, further explaining the different ratio of mature TgMIC5 to its pro-form in the Δ*cpc1* lysate. Although the maturation patterns of TgSUB1, TgM2AP, and TgMIC5 were changed in Δ*cpc1*, their subcellular trafficking patterns remained normal, suggesting that the extra amino acid residues associated with these micronemal effectors do not alter their subcellular targeting motifs. This mirrors a previous observation that the prepropeptide of TgSUB1 still results in the trafficking of a fused GFP to the microneme (40).

Among the three *Toxoplasma* cathepsin C-like orthologs, TgCPC1 showed the highest transcript abundance, followed by TgCPC2 (TGGT1_276130) (ToxoDB.org). TgCPC3 is speculated to be involved in oocyst development (17). The most similar ortholog of TgCPC2 within malaria parasites is PfDPAP3 (PF3D7_0404700). PfDPAP3, an essential gene in *Plasmodium spp*., was previously knocked down and identified as a key component mediating parasite invasion (21). The closest ortholog of TgCPC3 is the *Plasmodium* PfDPAP2 (PF3D7_1247800), which is a gametocyte-specific ortholog (22). The deletion of *PfDPAP2* causes the upregulation of PfDPAP1 by more than 2-fold, indicating a compensation mechanism between these two proteases. It is noteworthy that one chemical inhibitor of cathepsin C proteases, ML4118S, shows potency against parasites within both sexual and asexual stages (22). A previous report showed that TgCPC2 is localized to the PV and its mRNA level increased in the *TgCPC1*-deletion mutant (17). Our latest findings revealed that TgCPC2 is a rhoptry protein and that there was no altered regulation of mRNA between TgCPC1 and TgCPC2 within Δ*cpc1* (data not shown). Therefore, we speculate that TgCPC1 and TgCPC2 govern different subcellular events in *Toxoplasma* parasites.

Collectively, we characterized the roles of TgCPC1, a major *Toxoplasma* cathepsin C-like protease, in *Toxoplasma* infections. The ELC-localizing TgCPC1 plays an essential role in the activation of one subtilisin protease, TgSUB1 (**Fig. 8**). The defective TgSUB1 activation in Δ*cpc1* further results in the distribution of inactive TgSUB1 on the surface of the parasites that cannot properly trim a series of important invasion and egress effectors, including TgMIC2, TgAMA1, and TgPLP1. The absence of TgCPC1 leads to a significant loss of virulence in the parasites. A potent inhibitor against the malarial cathepsin C proteases did not show strong inhibition against TgCPC1. Future development of novel and specific inhibitors against *Toxoplasma* cathepsin C-like proteases can be utilized as a potential strategy for controlling toxoplasmosis.

**Figure 8.**
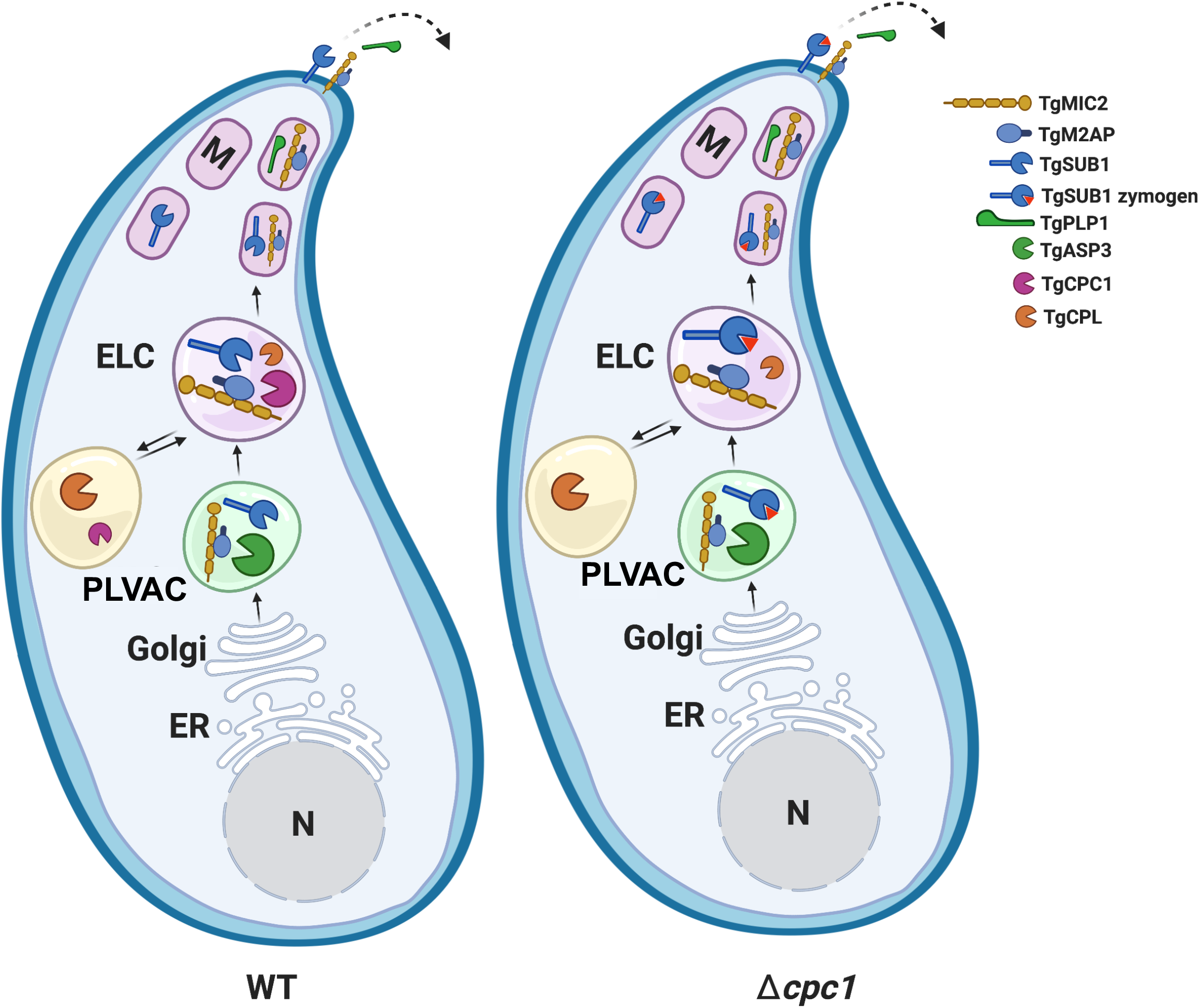
Working model of the post-translational modification of micronemal invasion effectors by TgCPC1 in *Toxoplasma*. Post ER biosynthesis, microneme proteins traffic through the Golgi apparatus and are cleaved within a post-Golgi compartment by TgASP3 (1), a major maturase for micronemal invasion effectors. Additionally, a minute amount of TgCPL makes an additional contribution to the maturation of some micronemal proteins in the ELC (2). Our findings suggest that TgCPC1, an aminopeptidase, conducts post-translational modification on some micronemal proteins before reaching their final forms, such as TgM2AP and TgAMA1, or performs initial trimming before subsequent cleavage, such as TgSUB1. Properly processed micronemal effectors are further delivered to microneme before subsequent processing on the parasite’s surface, followed by secretion. In the absence of TgCPC1, some incorrectly processed micronemal proteins are delivered to the surface and secreted from the parasites. Most importantly, TgSUB1 is kept as a zymogen on the parasite’s surface and it cannot cleave multiple key micronemal effectors required for parasite invasion and egress. ELC, endosome-like compartment; ER, endoplasmic reticulum; M, microneme; N, nucleus; PLVAC, plant-like vacuolar compartment.

## MATERIALS AND METHODS

### Ethical statement

This study was performed in compliance with the Public Health Service Policy on Humane Care and Use of Laboratory Animals and Association for the Assessment and Accreditation of Laboratory Animal Care guidelines. The animal protocol was approved by Clemson University’s Institutional Animal Care and Use Committee (Animal Welfare Assurance #: D16-00435; Protocol number: AUP2019-035). All efforts were made to minimize animal discomfort. CO_2_ overdose was used to euthanize mice. This form of euthanasia is consistent with the current recommendations of the Panel on Euthanasia of the American Veterinary Medical Association.

### Chemicals and reagents

All chemicals used in this study were ordered from VWR unless specified below. Zaprinast was acquired from Sigma Aldrich. The BI-2051 inhibitor was generously provided by opnMe.com. All PCR primers utilized in this study are listed in Table S1.

### Parasite culture

*Toxoplasma gondii* parasites were cultured at 37 °C with 5% CO_2_ within human foreskin fibroblast (HFF) cells (ATCC, SCRC-1041) or within hTERT cells in Dulbecco’s modified Eagle medium (DMEM), supplemented with 10% cosmic calf serum, 10 mM HEPES, pH 7.4, additional 2 mM _L_-glutamine, 1 mM pyruvate, and 100 U/mL penicillin/streptomycin. The parasites were purified by membrane filtration as previously described (50).

### Generation of transgenic parasites

The transgenic *Toxoplasma* strains generated and used in this work are listed in Table S2.

#### 1. 3xmyc-tagged TgCPC1 strains (TgCPC1-3xmyc^c^ and TgCPC1-3xmyc^i^)

In this study, a 3xmyc epitope was inserted at the C-terminus of TgCPC1 protein or engineered internally within a predicted antigenic region of the TgCPC1 protein (**Fig. 1A and Fig. S1B**). To generate the C-terminally 3xmyc-tagged TgCPC1 strain, a 3.9-kb DNA fragment upstream from the stop codon of TgCPC1 was amplified and cloned into p3xmyc-LIC-CAT plasmid. The resulting plasmid was linearized by BglII and introduced into RH Δ*ku80* parasites by electroporation. The 3xmyc tag was incorporated into the end of TgCPC1 gene by single crossover recombination. The resulting transfectants were selected by chloramphenicol and cloned out. The resulting strain was named TgCPC1-3xmyc^c^. To tag TgCPC1 internally with 3x epitope, the coding sequence of TgCPC1 was amplified from the parasite’s cDNA library by PCR and flanked with 1kb of its 5’- and 3’-UTRs using standard cloning techniques. The resulting TgCPC1 expression cassette was cloned into a plasmid vector carrying a bleomycin (BLE) resistance cassette to generate a wildtype TgCPC1 expression construct. Using NEB Q5-directed mutagenesis, the region encoding 3xmyc epitope was inserted to the expected location inside TgCPC1 indicated in Fig. S1B. The correct clone was verified by Sanger sequencing. Similarly, the resulting plasmid was electroporated into RH Δ*ku80* parasites, which was selected by bleomycin extracellularly twice prior to cloning. This internally 3xmyc-tagged strain was named TgCPC1-3xmyc^i^. Immunoblotting was used to confirm the expression of the 3xmyc-tagged TgCPC1 fusion proteins.

#### 2. *Tgcpc1*-null mutant (Δ*cpc1*) and the corresponding complementation strain (Δ*cpc1CPC1*)

To generate *Tgcpc1*-deficient parasites, 3 kb of the 5’ and 3’ UTR regions of the *TgCPC1* gene were PCR-amplified and flanked at both ends of a pyrimethamine resistance cassette (DHFR) to create a *Tgcpc1* deletion construct. RH Δ*ku80* parasites were electroporated with the *Tgcpc1* deletion construct, selected by pyrimethamine, and cloned out via limiting dilution. PCR and qPCR were used to confirm the successful ablation of*Tgcpc1* gene. To generate a TgCPC1 complementation strain, the *Tgcpc1*-deficient parasites were electroporated with the wildtype TgCPC1 expression construct mentioned above. The transfectants were selected by bleomycin at the extracellular stage and cloned out. PCR was used to confirm the integration of TgCPC1 into the parasite’s genome and qPCR was used to quantify the restored transcript level of TgCPC1.

### Transfection of parasites

Freshly lysed *Toxoplasma* parasites were syringed, filter-purified, and resuspended in Cytomix buffer (25 mM HEPES, pH 7.6, 120 mM KCl, 10 mM K_2_HPO4/ KH_2_PO4, 5 mM MgCl_2_, 0.15 mM CaCl_2_, and 2 mM EGTA). Parasites were pelleted and washed once in Cytomix buffer before they were resuspended at 2.5 × 10^7^ parasites per mL in Cytomix buffer. Four hundred microliters of the parasite resuspension was mixed with 20 µg DNA and 2 mM ATP/5 mM reduced glutathione in a total volume of 500 µL. The mixture was then electroporated at 2 kV and 50 ohm resistance using a BTX Gemini X2 (Harvard Apparatus). Next, the transfected parasites were transferred to an HFF-coated T25 flask and allowed to recover for 24 h prior to drug selection.

### Quantitative PCR (qPCR) assay

The WT, Δ*cpc1*, and Δ*cpc1CPC1* parasites were grown in HFF cells for 48 hrs and filter-purified for total RNA extraction using a Direct-zol RNA MiniPrep Plus kit (Zymo). Transcript levels of individual genes were determined by the Luna Universal One-Step RT-PCR kit (NEB) using approximately 100-200 ng of total RNA per sample as input. All qPCR assays were performed using the BioRad CFX96 Touch Real-Time PCR detection system. Data were analyzed by taking the cycle threshold (CT) values for each gene and using the double delta CT (ΔΔCT) analysis method to calculate the relative abundance of each target in the transgenic strains compared to WT control as described previously (50). *TgActin* was included as the housekeeping gene for normalization.

### Plaque assay

Freshly lysed parasites were purified as described above and resuspended in D10 medium at 100 tachyzoites per mL. Two hundred parasites were inoculated into individual wells of HFF-coated 6-well plates and allowed to grow for 7 days at 37 °C with 5% CO_2_. Post-incubation, medium was carefully aspirated to avoid disturbance of HFF monolayers and the plates were gently washed once with PBS, stained with 0.2% crystal violet for 5 min and de-stained with water until the plaques were clearly visualized. Plates were air-dried overnight, followed by phase-contrast imaging using a Leica DMi8 inverted epifluorescence microscope under 25x magnification. The number of plaques in each well were counted. At least 50 individual plaques were documented and their areas quantified by ImageJ as previously reported (51). Three biological replicates were combined for statistical significance calculation.

### Invasion assay

Freshly lysed parasites were syringed, filter purified, and resuspended at 5 × 10^7^ parasites per mL in invasion medium (DMEM supplemented with 3% cosmic calf serum). Two hundred microliters of the parasite resuspension was inoculated into each well of an 8-well chamber slide pre-seeded with HFF cells and parasites were allowed to invade host cells for 30 min before fixation with 4% formaldehyde for 20 min. Slides were immunostained with mouse anti-TgSAG1 monoclonal antibody (1:2000) for 1 h to label attached parasites followed by a secondary stain using goat anti-mouse IgG conjugated with Alexa 594 (red) (Invitrogen, 1:1000). Next, the slide was permeabilized with 0.1% Triton X-100 for 10 min, and then stained with a rabbit polyclonal anti-TgMIC5 antibody (1:1000) and goat anti-rabbit IgG conjugated with Alexa 488 (green) (Invitrogen, 1:1000) to label all parasites, including invaded and attached parasites. DAPI was also included for nuclear staining. Six fields of view for each strain were captured by a Leica DMi8 inverted epifluorescence microscope and ImageJ software was used for analysis. The following equation was used to calculate invasion efficiency of each strain: ([sum of green parasites] − [sum of red parasites]) ⁄ total host nuclei. The assay was repeated, at minimum, in three biological replicates.

### Replication assay

Freshly lysed parasites were filter-purified and used to inoculate individual wells of an 8-well chamber slide that was pre-seeded with HFF cells at approximately 1 × 10^5^ cells per well. Non-invaded parasites were washed off at 4 hrs post-inoculation. Invaded parasites were allowed to continue to replicate within host cells for an additional 24 hrs prior to fixation. Infected host cells were stained with a monoclonal anti-TgGRA7 antibody (1:1000) and DAPI for labeling individual parasitophorous vacuoles (PVs) and parasite nuclei, respectively. Stained parasites were observed and counted by immunofluorescence microscopy. One hundred PVs were enumerated for each strain and plotted based on the distribution of different-sized PVs. The average number of parasites per PV was calculated for comparison. The assay was performed in triplicate.

### Egress assay

Filter-purified tachyzoites were resuspended in D10 medium at 5 × 10^5^ parasites per mL. One hundred microliters of the parasite resuspension were inoculated into each well of a 96-well plate pre-seeded with confluent HFF cells. Parasites were allowed to replicate for 18–24 h prior to being washed and incubated in 50 µL of Ringer’s buffer (10 mM HEPES, pH 7.2, 3 mM NaH_2_PO_4_, 1 mM MgCl_2_, 2 mM CaCl_2_, 3 mM KCl, 115 mM NaCl, 10 mM glucose, and 1% FBS) for 20 min. Next, an equal volume of 1 mM zaprinast dissolved in Ringer’s buffer was added to all sample wells and incubated for 5 min at 37 °C and 5% CO_2_. The wells containing uninfected cells were treated with 50 µL of plain Ringer’s buffer or 1% Triton X-100 in Ringer’s buffer as negative and positive controls, respectively. Samples were spun at 1,000 *x g* for 5 min twice to pellet insoluble cell debris. The supernatant was collected and subjected to a standard lactate dehydrogenase release assay as previously described (50, 52). The assay was conducted in five independent replicates.

### Chemically induced motility analysis

35 mm MatTek dishes (MatTek Corporation) were treated with 10% fetal bovine serum (FBS) for 24 hrs before imaging to provide sufficient protein to allow a surface conducive for motility. Dishes were washed once with PBS and filled with 2 mL of Ringer without Ca^2+^ (pH 7.4), and then chilled on ice. Purified parasites were added to the dish and incubated on ice for 15 min. Non-attached parasites were removed by washing dishes with 2 mL of ice-cold Ringer’s buffer without Ca^2+^. Dishes were then transferred to the General Electric Delta Vision environmental chamber preset to 37°C and allowed to equilibrate temperature for 5 min. Time-lapse videos were recorded using an Olympus IX-71 inverted fluorescence microscope with a Photometrix CoolSnapHQ CCD camera driven by Delta Vision software. The exposure duration, gain, laser intensity, and filter settings were kept the same in all videos for quantification. After 30 sec, 100 mM zaprinast was added to dishes to stimulate motility. Tracings were measured via two different conditions: (A) To quantify circular motility, the total number of parasites in the field of view were divided by the total number of parasites completing at least one full circle movement. Data were derived from 6 independent trials. (B) For calculating the total distance traveled, ImageJ software with the MTrackJ plugin was used to track and calculate distance. Data were reported as the average distance traveled (in µm) of 4 parasites from 4 independent biological trials.

### Mouse studies

Six- to eight-week-old outbred CD-1 mice were infected by subcutaneous injection with 100 WT, Δ*cpc1*, and Δ*cpc1CPC1* parasites resuspended in PBS. Infected mice were monitored daily for symptoms for a 30-day period. Following the protocol approved by Clemson University’s IACUC, mice that appeared moribund were humanely euthanized via CO_2_ overdose. Enzyme-linked immunosorbent assay (ELISA) was used to check for seroconversion in the surviving mice. In addition, the survivors were allowed to rest for 10 days and challenged by subsequent infection with 1,000 WT parasites via subcutaneous inoculation to confirm previous infections. Mice were kept for an additional 30 days and monitored daily for symptoms.

### Immunofluorescence and co-localization assays

HFF cells were pre-seeded into an 8 well chamber slide and grown for 24 hrs prior to all assays. Freshly egressed parasites were used to infect chamber slides for either 30 min (pulse-invaded parasites) or 18-24 hrs (replicated parasites). To detect surface-localized TgSUB1, extracellular parasites were adhered to chamber slide wells prior to immunofluorescence assay. The immunofluorescence staining procedure was followed from a previous publication (50). A Leica DMi8 inverted fluorescent microscope equipped with a CCD camera was used to visualize and capture images. Image processing was completed using Leica LAS X software. Co-localization analysis of TgCPC1 with PLVAC or ELC was quantified by assessing the proximity between TgCPC1 with a PLVAC marker (TgCPL) or ELC markers (proTgM2AP and TgNHE3) within 75-80 parasites per strain. Data from four separate IFA experiments were compiled for plotting and statistical significance calculation by one-way ANOVA.

### Excretory secretory antigens (ESAs) preparation

Freshly lysed parasites were syringed, filter purified, and resuspended at 5 × 10^8^ parasites per mL in D1 medium (DMEM medium supplemented with 1% FBS). One hundred microliters of parasite resuspension were transferred to a microfuge tube and incubated at 37 °C for 30 min to prepare constitutive ESAs. Induced ESAs were obtained by treating the parasite resuspension with 1% (v/v) ethanol at 37 °C for 2 min. ESAs were separated from intact parasites by centrifugation at 1,000 x *g* for 10 min at 4 °C, then transferred to a new microfuge tube, mixed with SDS-PAGE sample loading buffer, and boiled for 5 min for downstream immunoblotting analysis.

### SDS-PAGE and Immunoblotting

Parasite lysates and ESA fractions were subjected to standard SDS-PAGE and immunoblotting procedures as described previously (50). In brief, based on the sizes of target proteins, samples were resolved on 7.5%, 10%, and 12% SDS-PAGE gels, and transferred to PVDF membranes using a semi-dry protein transfer system. Following transfer, 5% non-fat milk in PBS containing 0.1% Tween-20 (PBS-T buffer) was used as blocking buffer. Primary and secondary antibodies were diluted in 1% (w/v) non-fat milk in PBS-T at the titers reported before. SuperSignal WestPico chemiluminescent substrate (Thermo) was applied to the blots for the detection of target bands. The chemiluminescence signals were captured by Azure C600 Imaging System for documentation and further quantification by LI-COR Image Studio Lite.

### Estimation of apparent molecular weights of protein bands on SDS-PAGE

Individual TgCPC1-derived species were resolved by 12% SDS-PAGE from two independent trials and their relative distances (Rf) were measured by AzureSpot software (version 14.2). The Rf values of the protein standards with known molecular weights were also measured and plotted for creating a standard curve using the cubic spline curve algorithm to calculate the apparent molecular weights of cleaved TgCPC1 polypeptides.

### Molecular docking

The chemical structure of BI-2051 was drawn with ChemOffice professional 19 suite (PerkinElmer, Waltham, MA), and a three-dimensional (3D) structure was generated with VeraChem Vconf (VerChem LLC, Germantown, MD). The 3D structure was optimized by Gaussian 16 suite (Gaussian Inc., Wallingford, CT) with Density Functional Theory (DFT), employing the B3LYP/6-311G (d,p) level of theory (53). The 3D crystal structure of human dipeptidyl peptidase I (hDPP-I; PDB 2DJG) was retrieved from the RCSB protein data bank (26). The predicted 3D structures of *Plasmodium falciparum* dipeptidyl peptidase 1 (PfDPAP1) and *Toxoplasma gondii* cathepsin C-like proteases (TgCPC1) were retrieved from the AlphaFold protein structure database (43). The optimized BI-2051 and the proteins were prepared by removing co-crystallized ligands, heteroatoms, and water molecules, where applicable, using Pymol Molecular Graphics 2.0 (Schrödinger LLC, New York, NY), after which all structures were converted into pdbqt formats using AutoDock Tools (The Scripps Research Institute, La Jolla, CA). The coordinates of the active site residues of PfDPAP1 and TgCPC1 were aligned from the active site of hDPP-I based on conserved amino acid residues from BLASTp alignment between these homologs proteins. The BI-2051 was docked to the active sites of the proteins in vacuo using AutoDock vina with specific docking parameters and scoring functions described in the literature (54). The binding affinity of the ligand was measured in kcal/mol as a unit for a negative score (54). The binding conformation with the highest negative value was taken as the best pose for the corresponding protein-ligand complex. Subsequently, the best binding pose of each complex was analyzed using Pymol and Discovery Studio (Dassault Systémes, Waltham, MA) to reveal the protein-ligand interactions.

### Statistics

Prism software (GraphPad version 8) was used to perform statistical analysis for all data presented here. The specific statistical methods for each assay are specified within the figure legends.

## Acknowledgments

We thank our colleagues Drs. Michael Blackman, Peter Bradley, Vern Carruthers, Gary Ward, Jean Francois Dubremetz, and David Sibley for kindly providing key reagents for this study. We also want to thank Amy Bergmann for help with routine tissue culture preparation in the lab. This work was supported by National Institutes of Health grant R01AI143707 (to Z.D.), R01AI128356, and R01AI096836 (to S.N.J.M) and the grant from the Knights Templar Eye Foundation (to Z.D.). A.J.S. was partially funded by a UGA and T32 fellowship (T32AI060546). We declare that we have no conflicts of interest concerning the contents of this article.

## Supplemental Material

**Figure S1. Primary structure and motifs of TgCPC1**. (A) TgCPC1 carries a putative signal peptide. The prediction of the signal peptide was performed using a SignalP 6.0 algorithm (https://services.healthtech.dtu.dk/service.php?SignalP-6.0). (B) Antigenic region prediction was conducted using EMBOSS program for the internal epitope-tagging of TgCPC1. The region within the red box was picked as the site for insertion of the 3xmyc epitope tag. (C) Primary structure and motif annotation in TgCPC1-3xmyc^c^ and TgCPC1-3xmyc^i^ strains. The cleavage sites denoted by solid black arrowheads were deduced by comparing against cleavage sites within human CPC protease and the homologous alignment between TgCPC1 and human DPP-I. The cutting site between the putative light and heavy chains, indicated by the hollow black arrowheads, was predicted from the observed molecular weights of cleaved TgCPC1 species shown in Fig. 1A. The essential Cys, His, and Asn within the catalytic triad, are labeled in red. Asterisks represent the stop site of translation.

**Figure S2. TgCPC1 was not detected in the PV**. To test if TgCPC1 is secreted into the PV, the replicated TgCPC1-3xmyc^c^ and TgCPC1-3xmyc^i^ parasites were co-immunostained with anti-TgGRA7 and anti-myc antibodies. The myc staining was contained within the parasites and was not observed in the PV space denoted by arrowheads. Bar = 2 µm.

**Figure S3. A trace amount of TgCPC1 was secreted by *Toxoplasma* parasites**. Purified extracellular WT, TgCPC1-3xmyc^c^ and TgCPC1-3xmyc^i^ parasites were subjected to the preparation of constitutive ESAs. The ESAs were probed with anti-myc, anti-TgCPL (negative control), and anti-TgPI-1 (positive control) antibodies. In contrast to TgCPL staining, a trace amount of TgCPC1 was observed in the ESA fractions, suggesting that TgCPC1 can be released from the parasites by an undefined pathway. At least two independent preparations of ESAs and total protein lysates were generated for this assay.

**Figure S4. Generation of Δ*cpc1* and Δ*cpc1CPC1* strains**. (A) Schematic representation of the approach used for generating Δ*cpc1* and for complementing the parasites with *TgCPC1*. WT parasites were transfected with a deletion construct containing a DHFR resistance cassette flanked by the 5’ and 3’ UTR regions that are upstream and downstream of the *TgCPC1* gene. Homologous recombination allowed for the replacement of the *TgCPC1* gene with the DHFR resistance cassette in order to generate Δ*cpc1*. The Δ*cpc1* parasites were complemented by introducing a plasmid containing the coding sequence of *TgCPC1* flanked by its own 5’ and 3’ UTRs in addition to a bleomycin (*BLE*) resistance cassette. (B) PCR verification of Δ*cpc1* and Δ*cpc1CPC1* strains. The PCR primers indicated in the schematic were used to verify the absence and complementation of the *TgCPC1* coding sequence (CDS) within Δ*cpc1* and Δ*cpc1CPC1*, respectively. The sizes of the corresponding PCR products were indicated in the schematic. The band marked with asterisk was from non-specific PCR amplification. (C) Quantitative PCR confirmed the loss and recovery of *TgCPC1* transcripts in Δ*cpc1* and Δ*cpc1CPC1* parasites. *TgActin* was included as a loading control.

**Figure S5. The prediction of the active sites of PfDPAP1 and TgCPC1**. The protein sequences of hDPP-I, PfDPAP1, and TgCPC1 were acquired from www.unprot.org. A global BLASTp program was used for alignment. The amino acids in red are the conserved residues.

**Figure S6. TgCPL was not involved in the intracellular cleavage of TgCPC1 but affected the abundance of TgCPC1**. (A) TgCPC1 was tagged with C-terminal and internal 3xmyc tags in Δ*cpl*. WT, TgCPC1-3xmyc^c^, Δ*cpl*::*TgCPC1-3xmyc*^*c*^, TgCPC1-3xmyc^i^, and Δ*cpl::TgCPC1-3xmyc*^*i*^ parasites were grown in HFFs for 48 h before lysate preparation. Lysates were probed with anti-myc antibody to assess the cleavage patterns of TgCPC1. There were no distinguishable changes in TgCPC1 cleavage between WT and Δ*cpl* background, suggesting that TgCPL is not required for TgCPC1 proteolytic cleavage. TgCPL was also probed to confirm its loss in *TgCPL*-deletion strains. TgActin was included as a loading control. (B) To validate the observation shown in Fig. S6A, WT, TgCPC1-3xmyc^c^ and TgCPC1-3xmyc^i^ parasites were treated with 1 µM LHVS or DMSO (vehicle control) for 48 hrs before lysate preparation. Similar phenotypes were observed.

**Table S1. Primers used in the study**.

**Table S2. Parasite strains used in the study**.

## REFERENCES

1. Dogga SK, Mukherjee B, Jacot D, Kockmann T, Molino L, Hammoudi P-M, Hartkoorn RC, Hehl AB, Soldati-Favre D. 2017. A druggable secretory protein maturase of Toxoplasma essential for invasion and egress. Elife 6:223.

2. Parussini F, Coppens I, Shah PP, Diamond SL, Carruthers VB. 2010. Cathepsin L occupies a vacuolar compartment and is a protein maturase within the endo/exocytic system of Toxoplasma gondii. Mol Microbiol 76:1340–1357.

3. Blader IJ, Coleman BI, Chen C-T, Gubbels M-J. 2015. Lytic Cycle of Toxoplasma gondii: 15 Years Later. Annu Rev Microbiol 69:1–23.

4. Tenter AM, Heckeroth AR, Weiss LM. 2000. Toxoplasma gondii: from animals to humans. Int J Parasitol 30:1217–1258.

5. Lagal V, Binder EM, Huynh M, Kafsack BFC, Harris PK, Diez R, Chen D, Cole RN, Carruthers VB, Kim K. 2010. Toxoplasma gondii protease TgSUB1 is required for cell surface processing of micronemal adhesive complexes and efficient adhesion of tachyzoites. Cell Microbiol 12:1792–1808.

6. Harb OS, Roos DS. 2019. Toxoplasma gondii, Methods and Protocols. Methods Mol Biology 2071:27–47.

7. Stasic AJ, Moreno SNJ, Carruthers VB, Dou Z. 2022. The Toxoplasma plant-like vacuolar compartment (PLVAC). J Eukaryot Microbiol 69:e12951.

8. Miranda K, Pace DA, Cintron R, Rodrigues JCF, Fang J, Smith A, Rohloff P, Coelho E, Haas FD, Souza WD, Coppens I, Sibley LD, Moreno SNJ. 2010. Characterization of a novel organelle in Toxoplasma gondii with similar composition and function to the plant vacuole. Mol Microbiol 76:1358– 1375.

9. Dou Z, McGovern OL, Cristina MD, Carruthers VB. 2014. Toxoplasma gondii Ingests and Digests Host Cytosolic Proteins. Mbio 5:e01188–14.

10. McDonald C, Smith D, Cristina MD, Kannan G, Dou Z, Carruthers VB. 2020. Toxoplasma Cathepsin Protease B and Aspartyl Protease 1 Are Dispensable for Endolysosomal Protein Digestion. Msphere 5:e00869–19.

11. Dou Z, Coppens I, Carruthers VB. 2013. Non-canonical maturation of two papain-family proteases in Toxoplasma gondii. J Biol Chem 288:3523–3534.

12. Cristina MD, Dou Z, Lunghi M, Kannan G, Huynh M-H, McGovern OL, Schultz TL, Schultz AJ, Miller AJ, Hayes BM, Linden W van der, Emiliani C, Bogyo M, Besteiro S, Coppens I, Carruthers VB. 2017. Toxoplasma depends on lysosomal consumption of autophagosomes for persistent infection. Nat Microbiol 2:17096.

13. Liu J, Pace D, Dou Z, King TP, Guidot D, Li Z, Carruthers VB, Moreno SNJ. 2014. A vacuolar-H+-pyrophosphatase (TgVP1) is required for microneme secretion, host cell invasion, and extracellular survival of Toxoplasma gondii. Mol Microbiol 93:698–712.

14. Stasic AJ, Chasen NM, Dykes EJ, Vella SA, Asady B, Starai VJ, Moreno SNJ. 2019. The Toxoplasma Vacuolar H+-ATPase Regulates Intracellular pH and Impacts the Maturation of Essential Secretory Proteins. Cell Reports 27:2132-2146.e7.

15. Turk B, Turk D, Turk V. 2000. Lysosomal cysteine proteases: more than scavengers. Biochimica Et Biophysica Acta Bba - Protein Struct Mol Enzym 1477:98–111.

16. Korkmaz B, Caughey GH, Chapple I, Gauthier F, Hirschfeld J, Jenne DE, Kettritz R, Lalmanach G, Lamort A-S, Lauritzen C, Legowska M, Lesner A, Marchand-Adam S, McKaig SJ, Moss C, Pedersen J, Roberts H, Schreiber A, Seren S, Thakkar NS. 2018. Therapeutic targeting of cathepsin C: from pathophysiology to treatment. Pharmacol Therapeut 190:202–236.

17. Que X, Engel JC, Ferguson D, Wunderlich A, Tomavo S, Reed SL. 2007. Cathepsin Cs are key for the intracellular survival of the protozoan parasite, Toxoplasma gondii. J Biol Chem 282:4994–5003.

18. Rosenthal PJ. 2004. Cysteine proteases of malaria parasites. Int J Parasitol 34:1489–1499.

19. Rosenthal PJ, Sijwali PS, Singh A, Shenai BR. 2002. Cysteine proteases of malaria parasites: targets for chemotherapy. Curr Pharm Design 8:1659–72.

20. Klemba M, Gluzman I, Goldberg DE. 2004. A Plasmodium falciparum dipeptidyl aminopeptidase I participates in vacuolar hemoglobin degradation. J Biol Chem 279:43000–43007.

21. Lehmann C, Tan MSY, Vries LE de, Russo I, Sanchez MI, Goldberg DE, Deu E. 2018. Plasmodium falciparum dipeptidyl aminopeptidase 3 activity is important for efficient erythrocyte invasion by the malaria parasite. Plos Pathog 14:e1007031.

22. Tanaka TQ, Deu E, Molina-Cruz A, Ashburne MJ, Ali O, Suri A, Kortagere S, Bogyo M, Williamson KC. 2013. Plasmodium Dipeptidyl Aminopeptidases as Malaria Transmission-Blocking Drug Targets. Antimicrob Agents Ch 57:4645–4652.

23. Barylyuk K, Koreny L, Ke H, Butterworth S, Crook OM, Lassadi I, Gupta V, Tromer E, Mourier T, Stevens TJ, Breckels LM, Pain A, Lilley KS, Waller RF. 2020. A Comprehensive Subcellular Atlas of the Toxoplasma Proteome via hyperLOPIT Provides Spatial Context for Protein Functions. Cell Host Microbe 28:752-766.e9.

24. Bohley P, Seglen PO. 1992. Proteases and proteolysis in the lysosome. Experientia 48:151–157.

25. Turk D, Janjić V, Stern I, Podobnik M, Lamba D, Dahl SW, Lauritzen C, Pedersen J, Turk V, Turk B. 2001. Structure of human dipeptidyl peptidase I (cathepsin C): exclusion domain added to an endopeptidase framework creates the machine for activation of granular serine proteases. Embo J 20:6570–6582.

26. Mølgaard A, Arnau J, Lauritzen C, Larsen S, Petersen G, Pedersen J. 2006. The crystal structure of human dipeptidyl peptidase I (cathepsin C) in complex with the inhibitor Gly-Phe-CHN2. Biochem J 401:645–50.

27. Pszenny V, Davis PH, Zhou XW, Hunter CA, Carruthers VB, Roos DS. 2012. Targeted Disruption of Toxoplasma gondii Serine Protease Inhibitor 1 Increases Bradyzoite Cyst Formation In Vitro and Parasite Tissue Burden in Mice. Infect Immun 80:1156–1165.

28. Huynh M-H, Carruthers VB. 2006. Toxoplasma MIC2 is a major determinant of invasion and virulence. Plos Pathog 2:e84.

29. Huynh M-H, Rabenau KE, Harper JM, Beatty WL, Sibley LD, Carruthers VB. 2003. Rapid invasion of host cells by Toxoplasma requires secretion of the MIC2-M2AP adhesive protein complex. Embo J 22:2082–2090.

30. Jewett TJ, Sibley LD. 2004. The Toxoplasma Proteins MIC2 and M2AP Form a Hexameric Complex Necessary for Intracellular Survival. J Biol Chem 279:9362–9369.

31. Kafsack BFC, Pena JDO, Coppens I, Ravindran S, Boothroyd JC, Carruthers VB. 2009. Rapid membrane disruption by a perforin-like protein facilitates parasite exit from host cells. Science 323:530–533.

32. Parussini F, Tang Q, Moin SM, Mital J, Urban S, Ward GE. 2012. Intramembrane proteolysis of Toxoplasma apical membrane antigen 1 facilitates host-cell invasion but is dispensable for replication. Proc National Acad Sci 109:7463–7468.

33. Santos JM, Ferguson DJP, Blackman MJ, Soldati-Favre D. 2011. Intramembrane cleavage of AMA1 triggers Toxoplasma to switch from an invasive to a replicative mode. Science 331:473–477.

34. Roiko MS, Carruthers VB. 2013. Functional Dissection of Toxoplasma gondii Perforin-like Protein 1 Reveals a Dual Domain Mode of Membrane Binding for Cytolysis and Parasite Egress. J Biol Chem 288:8712–8725.

35. Harper JM. 2006. A Cleavable Propeptide Influences Toxoplasma Infection by Facilitating the Trafficking and Secretion of the TgMIC2-M2AP Invasion Complex. Mol Biol Cell 17:4551–4563.

36. Brydges SD, Harper JM, Parussini F, Coppens I, Carruthers VB. 2008. A transient forward-targeting element for microneme-regulated secretion in Toxoplasma gondii. Biol Cell 100:253–264.

37. Morlon-Guyot J, Hajj HE, Martin K, Fois A, Carrillo A, Berry L, Burchmore R, Meissner M, Lebrun M, Daher W. 2018. A proteomic analysis unravels novel CORVET and HOPS proteins involved in Toxoplasma gondii secretory organelles biogenesis. Cell Microbiol 20:e12870.

38. Sloves P-J, Delhaye S, Mouveaux T, Werkmeister E, Slomianny C, Hovasse A, Alayi TD, Callebaut I, Gaji RY, Schaeffer-Reiss C, Dorsselear AV, Carruthers VB, Tomavo S. 2012. Toxoplasma sortilin-like receptor regulates protein transport and is essential for apical secretory organelle biogenesis and host infection. Cell Host Microbe 11:515–527.

39. Miller SA, Binder EM, Blackman MJ, Carruthers VB, Kim K. 2001. A Conserved Subtilisin-like Protein TgSUB1 in Microneme Organelles of Toxoplasma gondii. J Biol Chem 276:45341–45348.

40. Binder EM, Lagal V, Kim K. 2008. The Prodomain of Toxoplasma gondii GPI-Anchored Subtilase TgSUB1 Mediates its Targeting to Micronemes. Traffic 9:1485–1496.

41. Sanchez MI, Vries LE de, Lehmann C, Lee JT, Ang KK, Wilson C, Chen S, Arkin MR, Bogyo M, Deu E. 2019. Identification of Plasmodium dipeptidyl aminopeptidase allosteric inhibitors by high throughput screening. Plos One 14:e0226270.

42. Deu E, Leyva MJ, Albrow VE, Rice MJ, Ellman JA, Bogyo M. 2010. Functional Studies of Plasmodium falciparum Dipeptidyl Aminopeptidase I Using Small Molecule Inhibitors and Active Site Probes. Chem Biol 17:808–819.

43. Jumper J, Evans R, Pritzel A, Green T, Figurnov M, Ronneberger O, Tunyasuvunakool K, Bates R, Žídek A, Potapenko A, Bridgland A, Meyer C, Kohl SAA, Ballard AJ, Cowie A, Romera-Paredes B, Nikolov S, Jain R, Adler J, Back T, Petersen S, Reiman D, Clancy E, Zielinski M, Steinegger M, Pacholska M, Berghammer T, Bodenstein S, Silver D, Vinyals O, Senior AW, Kavukcuoglu K, Kohli P, Hassabis D. 2021. Highly accurate protein structure prediction with AlphaFold. Nature 596:583–589.

44. Liu Y, Zou X, Ou M, Ye X, Zhang B, Wu T, Dong S, Chen X, Liu H, Zheng Z, Zhao J, Wu J, Liu D, Wen Z, Wang Y, Zheng S, Zhu K, Huang X, Du X, Liang J, Luo X, Xie Y, Wu M, Lu C, Xie X, Liu K, Yuting Y, Qi G, Jing C, Yang G. 2019. Toxoplasma gondii Cathepsin C1 inhibits NF-κB signalling through the positive regulation of the HIF-1α/EPO axis. Acta Tropica: 195:35–43.

45. Nelson DC, Garbe J, Collin M. 2011. Cysteine proteinase SpeB from Streptococcus pyogenes – a potent modifier of immunologically important host and bacterial proteins. Biol Chem 392:1077–1088.

46. Que X, Reed SL. 1997. The role of extracellular cysteine proteinases in pathogenesis of Entamoeba histolytica invasion. Parasitol Today 13:190–194.

47. Dahl SW, Halkier T, Lauritzen C, Dolenc I, Pedersen J, Turk and V, Turk B. 2001. Human Recombinant Pro-dipeptidyl Peptidase I (Cathepsin C) Can Be Activated by Cathepsins L and S but Not by Autocatalytic Processing. Biochemistry 40:1671–1678.

48. Larson ET, Parussini F, Huynh M-H, Giebel JD, Kelley AM, Zhang L, Bogyo M, Merritt EA, Carruthers VB. 2009. Toxoplasma gondii cathepsin L is the primary target of the invasion-inhibitory compound morpholinurea-leucyl-homophenyl-vinyl sulfone phenyl. J Biol Chem 284:26839–26850.

49. Saouros S, Dou Z, Henry M, Marchant J, Carruthers VB, Matthews S. 2012. Microneme Protein 5 Regulates the Activity of Toxoplasma Subtilisin 1 by Mimicking a Subtilisin Prodomain. J Biol Chem 287:36029–36040.

50. Thornton LB, Teehan P, Floyd K, Cochrane C, Bergmann A, Riegel B, Stasic AJ, Cristina MD, Moreno SNJ, Roepe PD, Dou Z. 2019. An ortholog of Plasmodium falciparum chloroquine resistance transporter (PfCRT) plays a key role in maintaining the integrity of the endolysosomal system in Toxoplasma gondii to facilitate host invasion. Plos Pathog 15:e1007775.

51. Bergmann A, Floyd K, Key M, Dameron C, Rees KC, Thornton LB, Whitehead DC, Hamza I, Dou Z. 2020. Toxoplasma gondii requires its plant-like heme biosynthesis pathway for infection. PLoS pathogens 16:e1008499.

52. Kaja S, Payne AJ, Singh T, Ghuman JK, Sieck EG, Koulen P. 2015. An optimized lactate dehydrogenase release assay for screening of drug candidates in neuroscience. J Pharmacol Toxicol 73:1–6.

53. Lee C, Yang W, Parr RG. 1988. Development of the Colle-Salvetti correlation-energy formula into a functional of the electron density. Phys Rev B 37:785–789.

54. Trott O, Olson AJ. 2010. AutoDock Vina: Improving the speed and accuracy of docking with a new scoring function, efficient optimization, and multithreading. J Comput Chem 31:455–461.

